# Systems-level imaging reveals assembly and synaptic delivery of a multi-organelle ensemble during T cell killing

**DOI:** 10.64898/2026.10.30.763224

**Authors:** Akaash Kumar, Jane C. Stinchcombe, Gillian M. Griffiths

**Author notes:** Correspondence &.

## Abstract

Cytotoxic T lymphocytes (CTLs) undergo a dynamic intracellular polarisation of their secretory compartments as they recognise and destroy cancerous cells. How multiple organelles are coordinated in space and time during this process is unknown. To enable high-dimensional systems-level imaging of multiple organelles and their relationships at high spatiotemporal resolution we use multispectral super-resolution microscopy and volumetric live-cell imaging of CTLs interacting with their targets. Organelle polarisation is initiated by perinuclear endoplasmic reticulum (ER) contacting the immune synapse, while calcium signalling promotes multi-organelle clustering around the centrosome at the rear of the cell. This creates a multi-organelle ensemble that then translocates around the nucleus to the synapse. We track organelle interactions during dynamic polarisation revealing a shift from ER-dominated to Golgi-dominated contacts as organelles reach the synapse. These findings show that CTL polarity is built through staged, calcium-driven formation of a multi-organelle assembly. More broadly, this work establishes a framework for systems-level imaging of the organelle network in cells as they deliver effector functions.

## Introduction

Cytotoxic T lymphocytes (CTLs) kill infected and malignant cells by focusing secretion to a specialised immune synapse (IS) formed with the target cell [1–3]. This response is organised around the polarisation of the centrosome and secretory machinery towards the target [2, 4]. The centrosome docks at the synaptic plasma membrane thereby positioning the minus ends of the microtubule network at the site of secretion, allowing specialised secretory lysosomes termed cytotoxic granules to be delivered to the synapse [2, 3, 5, 6]. Much has been learned about the temporal order of events during polarisation including how synaptic actin dynamics control secretion [7–9] and nuclear polarisation facilitates an early transcriptional response [10, 11]. However, insights have been limited by the number of organelles that can be imaged simultaneously.

Multiple organelles are implicated in T cell activation. For instance, the ER Ca^2+^ sensor STIM1 and the plasma membrane channel ORAI1 accumulate at the T cell-target interface, and both synaptic puncta and distal STIM1-ORAI1 caps have been observed during T cell activation [12–14]. Additionally, mitochondria contribute to Ca^2+^ buffering and sustained Ca^2+^ entry [14, 15]. These observations suggest that the intracellular response downstream of TCR engagement may extend beyond reorientation of the centrosome and lytic granules to involve coordinated reorganisation of a wider organelle network. Yet how organelles interact with one another in CTLs has not been mapped, and no study has followed multiple organelles simultaneously in the same CTL during killing. Whether organelles reach the immune synapse as independent cargoes or as a coordinated architecture therefore remains unknown.

Membrane contact sites provide one mechanism through which such coordination could occur. At these interfaces, distinct organelle membranes are closely apposed without fusing, which enables the exchange of lipids, ions and other signalling molecules, coupling organelle dynamics and metabolism [16–19]. For instance, at ER-PM contacts, ER-resident STIM1 activated by ER Ca^2+^ depletion binds to ORAI1 channels in the plasma membrane to drive store-operated calcium entry (SOCE) [20–23]. The ER also forms functionally important contact sites with mitochondria, endolysosomal compartments and the Golgi to regulate organelle fission, fusion and lipid transfer [18, 19]. Yet, like the cytotoxic response itself, organelle contact sites have generally been studied one pair at a time and have not been globally studied in the context of CTL killing.

Systems-level spectral imaging in COS-7 cells revealed a dynamic organelle interactome containing higher-order proximities among multiple organelles [24], while multispectral analysis of activated macrophages identified functional multi-organelle units that support inflammatory metabolism [25]. Whether an acute cellular response such as cytotoxicity rewires the CTL organelle interactome, and how such a network moves in space and time during killing events, is unknown.

Resolving the intracellular organisation of CTLs during killing requires operating in two measurement scales that are difficult to reconcile. Membrane contact sites are appositions of tens of nanometres, often within crowded regions of the cell. Identifying them therefore requires imaging beyond the diffraction limit and, ultimately, ultrastructural confirmation. The cytotoxic response, by contrast, unfolds over seconds to minutes across the full three-dimensional cell volume and requires many organelles to be followed simultaneously through spectral multiplexing, which is beyond the reach of conventional technologies. Sequential emission-based imaging becomes progressively slower as spectral channels are added, while spectral unmixing performs poorly in the low-signal-to-noise-ratio (SNR) environment typical of live-cell microscopy.

Here we address these technological constraints by deploying multispectral imaging frameworks to operate at different spatiotemporal scales. To characterise organelle interactions and contact sites at high spatial resolution, we develop excitation-multiplexed Airyscan super-resolution imaging. This resolves six organelle markers in single cells at 120 nm lateral resolution, and we validate the contacts it reveals in three-dimensional electron tomography. To follow the reorganisation of the organelle network over time, we apply our recently-established eight-camera multispectral imaging module with Richardson-Lucy Spectral Unmixing (RLSU) [26], which captures eight spectral channels simultaneously and recovers low-SNR live-imaging data. We deploy multi-spectral live-cell imaging on both spinning-disk confocal and oblique-plane light-sheet microscopes, combined with volumetric object analysis.

We find that organelle polarisation to the immune synapse follows a characteristic three-phase programme. The perinuclear ER first polarises towards the target; multiple organelles subsequently cluster around the centrosome at the rear of the cell regulated by calcium fluxes; and the resulting architecture is then delivered to the immune synapse. We call this process **c**entrosome **o**rganised **m**ulti-organelle **e**nsemble **t**rafficking (COMET). This reorganisation is accompanied by selective remodelling of organelle contacts, which are established away from the synapse and subsequently redistribute toward it. Together, these findings establish a systems-level view of CTL intracellular organisation in which cytotoxicity involves the coordinated assembly, remodelling and polarised delivery of a multi-organelle ensemble, and establish the CTL organelle interactome as a measurable cellular state.

## Results

### Excitation-multiplexed super-resolution microscopy enables systems-level imaging below the diffraction limit

To map organelle organisation at the systems-level, we developed an excitation-based multispectral imaging framework that extends Airyscan super-resolution microscopy beyond conventional low-plex imaging. Existing approaches impose a trade-off between spectral dimensionality and spatial resolution. Conventional multispectral microscopy typically separates fluorophores from their emission spectra using point-scanning confocal microscopes equipped with multi-channel spectral detectors (*e*.*g*., the Zeiss Quasar detector), providing spectral flexibility but remaining constrained by diffraction-limited confocal resolution. Conversely, Airyscan detection improves spatial resolution and is readily implemented on commercial microscopes, but has largely been used for two- or three-colour imaging, limiting its ability to resolve crowded intracellular systems containing many simultaneously labelled structures.

We therefore developed an excitation-multiplexed Airyscan acquisition strategy in which complete three-dimensional volumes were recorded sequentially under six excitation wavelengths on a Zeiss LSM 980 Airyscan. As commonly used fluorophores exhibit broad and overlapping excitation and emission spectra, each recorded volume contained contributions from multiple labelled structures. We computationally separated these mixed signals using our Richardson-Lucy Spectral Unmixing (RLSU) algorithm [26]. RLSU uses an iterative, Poisson-aware reconstruction to reassign spectrally overlapping fluorescence to its underlying source components and outperforms linear unmixing, HyU and non-negative matrix factorisation, particularly at the low signal-to-noise ratios encountered in fluorescence imaging.

This combination of excitation multiplexing, Airyscan detection and RLSU enables spectral multiplexing below the diffraction-limit. It allows six labelled cellular components to be resolved within the same cell while retaining the enhanced spatial resolution of Airyscan imaging, with measured lateral full-width-at-half-maximum (FWHM) values of 120-200 nm compared with 250-400 nm for conventional confocal acquisition. The resulting framework provides a route to systems-level visualisation of densely packed intracellular structures at spatial scales that are inaccessible to conventional multi-spectral confocal microscopy.

We next tested whether the extensive spectral overlap inherent to this acquisition strategy could be accurately unmixed. In simulations, six spatially distinct ground-truth objects were assigned six different fluorophores (Supplementary Figure S1a-c), and a forward model was constructed incorporating the excitation laser lines, dichroic beam splitters, emission filters and fluorophore photophysical properties (Supplementary Figure S1d-f). These ground-truth objects were projected through the resulting mixing matrix to generate spectrally mixed images, to which Poisson noise was added to reproduce photon-limited fluorescence measurements (Supplementary Figure S1g). Application of RLSU recovered the individual underlying structures from these mixed data (Supplementary Figure S1h). The six-fluorophore mixing matrix was well conditioned (condition number, 3.16), and reconstruction yielded a maximum root-mean-square error (RMSE) of 1.13 across the recovered channels. Thus, despite substantial spectral overlap, excitation-multiplexed Airyscan imaging combined with RLSU can accurately recover multiple labelled structures while preserving the spatial resolution advantage of Airyscan detection.

### Target recognition redistributes a broad organelle network to the synapse

We first asked whether CTLs undergo a global change in intracellular organisation during the transition from migration to killing. Primary OT-I cells were labelled simultaneously for multiple organelles using a combination of polycistronic plasmids and live-cell dyes, and imaged by our multispectral Airyscan microscopy framework during migration and after engagement with OVA-pulsed EL4 target cells (Figure 1a, Supplementary Figure S2).

**Figure 1:**
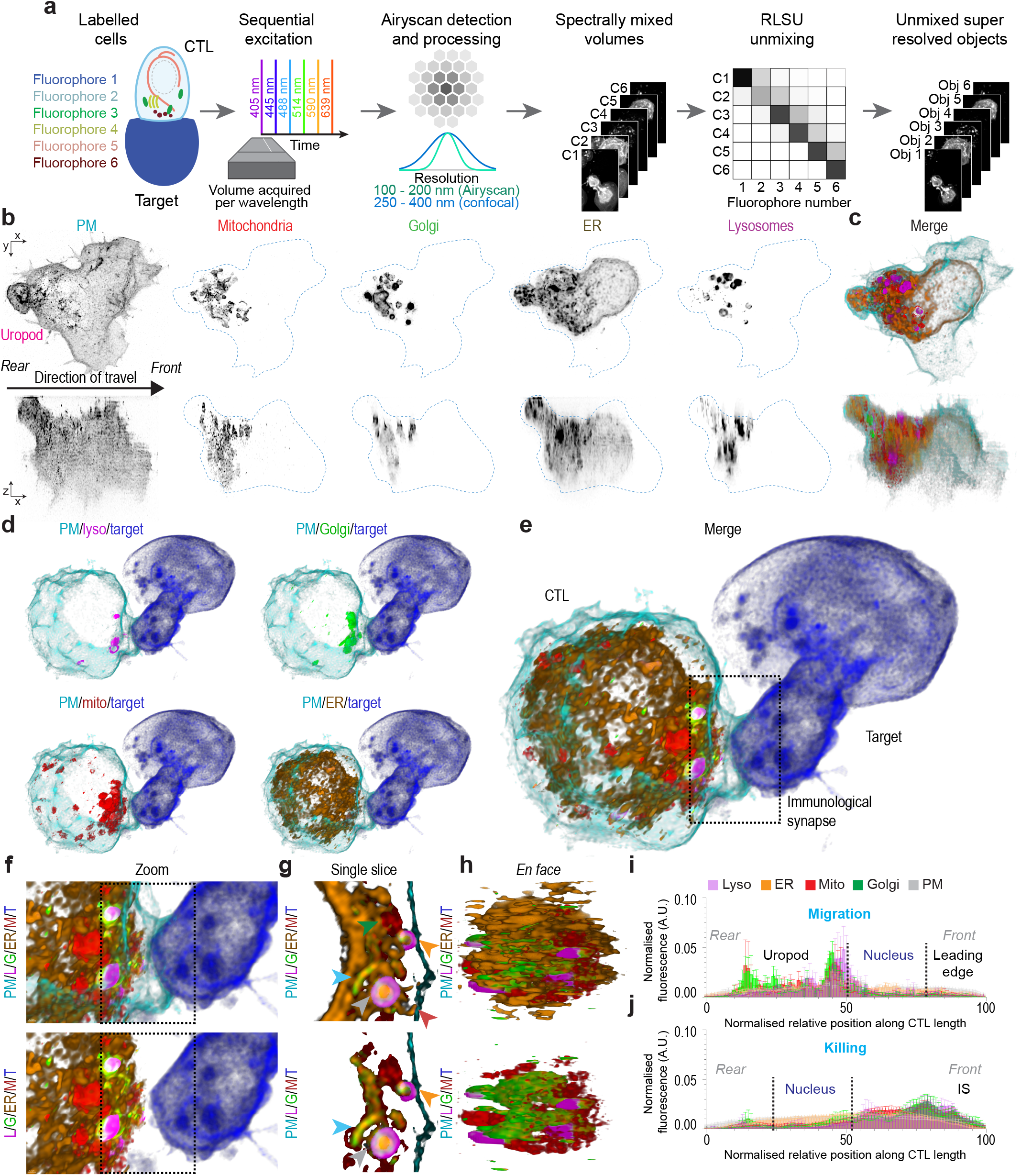
Excitation-multiplexed Airyscan imaging reveals broad organelle redistribution during CTL killing. **a** Schematic of multispectral Airyscan imaging and RLSU processing of CTLs. **b, c** Individual channels and merged views of a representative migrating CTL (*n* = 6 cells, *N* = 3 independent experiments), shown as xy and xz projections. Dashed cyan line indicates CTL boundary. **d, e** Corresponding views of a representative target-engaged CTL (*n* = 6 cells, *N* = 3 independent experiments). Target cell is shown in blue. Dotted box indicates the immune synapse. **f** Enlarged view of the dotted box shown in (e) indicating the immune synapse, with plasma membrane signals omitted in the bottom panel. **g** Single slice of the immune synapse shown in (e) as a colour merge of all six channels. ER channel is omitted in the bottom panel. Arrows indicate regions of proximity between various organelles (red: ER-PM, green: mitochondria-ER, grey: mitochondria-Golgi-lysosome-ER, orange: lysosome-ER-PM, cyan: mitochondria-Golgi-ER). **h** *En face* views of the immune synapse. ER channel is omitted in the bottom panel. **i, j** Normalised fluorescence profiles of organelles along the CTL rear-to-front axis during migration and target engagement, respectively (*n* = 6 cells per state, *N* = 3 independent experiments). PM, plasma membrane; ER, endoplasmic reticulum; M, Mito, mitochondria; L, Lyso, lysosomes; G, Golgi; T, target. Display colours follow the panel keys.

Migrating CTLs displayed a highly polarised intracellular organisation. The Golgi apparatus, mitochondria, lysosomes and the peripheral ER were concentrated at the rear of the cell behind the nucleus, within or adjacent to the uropod. Fewer than 5% of the segmented organelle volumes were positioned ahead of the nucleus in the analysed cells (Figure 1b-c,i, Supplementary Video 1).

Target cell engagement reversed this polarity. Within approximately 10 min, the uropod was lost and ER, Golgi, mitochondria and lysosomes became tightly polarised to the immune synapse (Figure 1d-f, Supplementary Video 2). *En face* views of the synapse revealed a spatially ordered organelle arrangement with the Golgi centrally positioned surrounded by mitochondria, lysosomes and ER tubules (Figure 1h). This reorganisation highlights the rapid and coordinated redistribution of organelles during the transition from migration to target cell killing and extends the classical reorientation of the centrosome and secretory lysosomes [2, 4–6] to a broader set of intracellular compartments.

The multispectral images also revealed extensive sub-resolution apposition among organelles at the synapse, including ER-PM, ER-mitochondria, ER-lysosome, Golgi-ER, Golgi-lysosome, mitochondria-lysosome and mitochondria-Golgi relationships, as well as regions in which three or more labelled compartments converged (Figure 1g). Target recognition therefore appears to alter not only the position of individual organelles but also the spatial relationships between them. However, as light microscopy does not resolve the membrane-to-membrane distances required to define *bona fide* membrane contact sites [16], we refer to these relationships simply as organelle contacts unless independently validated ultrastructurally.

### The immune synapse contains extensive membrane contact sites between polarised organelles

We next asked whether the polarised organelles form a defined network of organelle contacts at the immune synapse. To quantify this, we developed an analytical pipeline which segments the labelled organelles in migrating CTLs (Figure 2a) and CTL-target conjugates (Figure 2b) and quantifies shared voxels between 3D segmented objects at distances below the optical point-spread function of the microscope. During killing, contacts among multiple organelle pairs increased and became concentrated at the immune synapse, including lysosome–ER, Golgi–lysosome, mitochondria–lysosome, ER–mitochondria, mitochondria–Golgi and plasma membrane–ER relationships (Figure 2c-j). The resulting pattern was not dominated by a single organelle pair, indicating broad restructuring of the organelle interactome.

**Figure 2:**
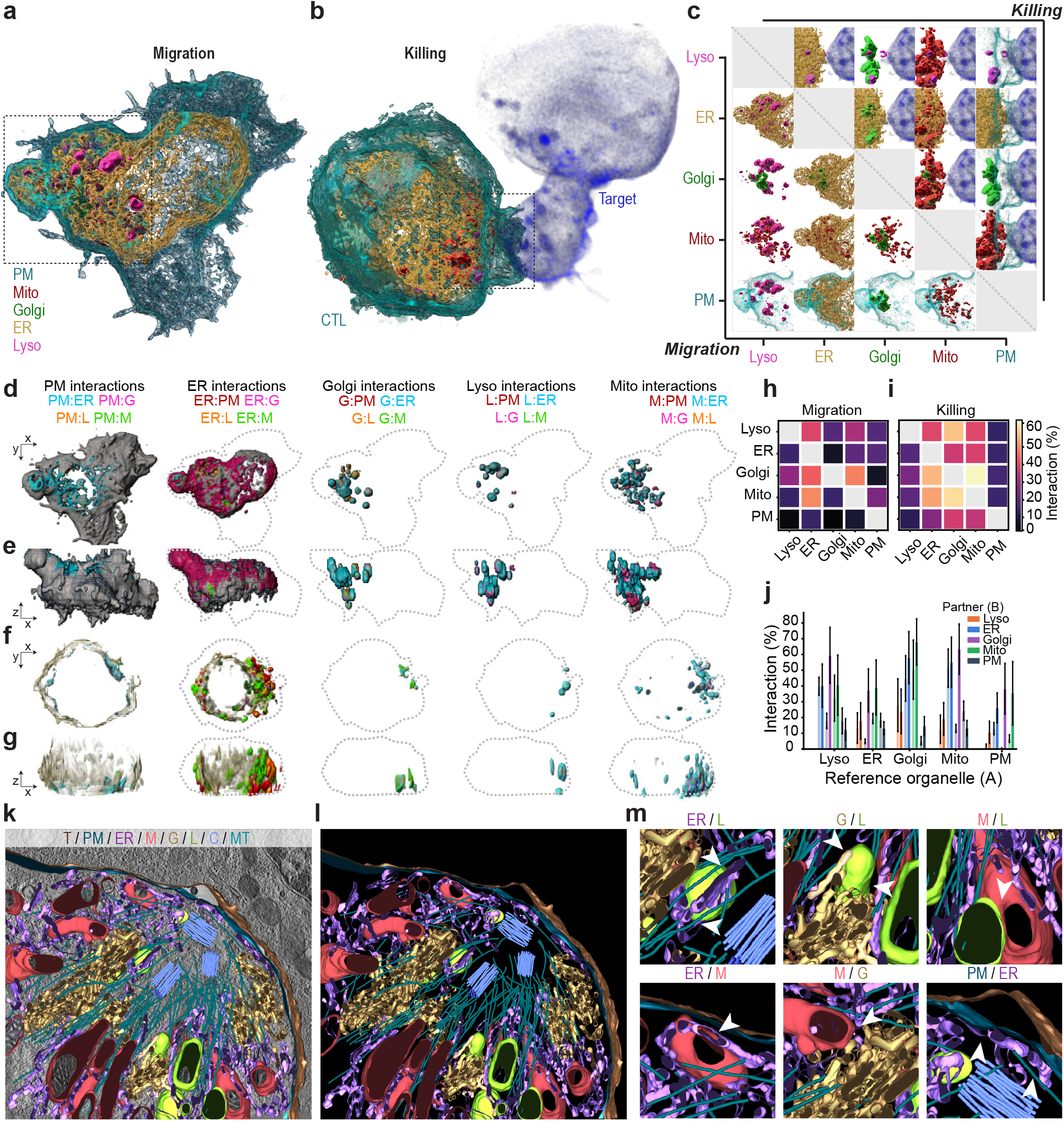
Organelle contact mapping and ultrastructural validation of membrane contact sites at the immune synapse. **a-b** Colour merge of 3D segmented organelles in a migrating CTL **a** or during killing **b. c** Pairwise organelle contacts between segmented organelles within the dashed box in **a** and **b**. Lower half of pairwise organelle contact matrix depicts organelle contacts in a migrating CTL, upper half depicts organelle contacts in a CTL during killing. **d-g** Colour-coded organelle interaction maps in a representative migrating CTL **d-e** and a representative CTL during killing **f-g** (*n* = 6 cells per state, *N* = 3 independent experiments). Top down (d,f) and orthogonal views (e,g) shown. Grey, underlying organelle mask; blue, interactions involving the ER; red, interactions involving the PM; magenta, interactions involving the Golgi; orange, interactions involving lysosomes; green, interactions involving mitochondria. Dashed line indicates cell boundaries. **h-i** Pairwise contact matrix of segmented voxels for each organelle interfacing every other organelle (contact distance below the point spread function of the microscope) displayed as a percentage for CTLs in their migratory state **h** or during killing **i** (mean shown, *n* = 6 cells per state, *N* = 3 independent experiments). **j** Percentage of voxels shared between reference organelle (A) and its partner (B) (mean and SEM shown, *n* = 6 cells per state, *N* = 3 independent experiments). Lighter coloured bars represent data deriving from CTLs in their migratory state whilst darker coloured bars represent data deriving from CTLs during killing. **k** 3D electron tomogram of organelles at the immune synapse. Raw data and segmentations of each organelle shown. **l** 3D electron tomogram of organelles at the immune synapse (3D segmentations of organelles shown without raw data). **m** Insets from panel (l) demonstrating various membrane contact sites between organelles at the immune synapse. White arrows indicate membrane contact sites between described organelle pairs. For (k-m): PM, plasma membrane (teal); T, target cell plasma membrane (brown); ER, endoplasmic reticulum (purple); M, mitochondria (red); G, Golgi apparatus (gold); L, lysosomes (green); C, centrosomes (blue); MT, microtubules (indigo).

To determine whether the close appositions identified by our multispectral super-resolution microscopy pipeline corresponded to *bona fide* membrane contact sites, we performed three-dimensional electron tomography of the immune synapse (Supplementary Video 3). Tomographic reconstructions revealed polarised centrioles flanked by dense ER sheets and tubules and surrounded by Golgi, mitochondria and lysosomal/lytic granule compartments (Figure 2k,l). Numerous regions of parallel membrane apposition of less than 30 nm were visible, including ER-PM, ER-mitochondria, ER-lytic-granule, Golgi-mitochondria and mitochondria-lytic-granule membrane contact sites, validating the light microscopy data at nanometre resolution (Figure 2m). These ultrastructural data establish that the immune synapse is not simply crowded with independently polarised organelles but it contains a dense network of *bona fide* inter-organelle membrane contact sites.

The differences between migrating and target-engaged CTLs indicate organelle remodelling during cytotoxic responses but they do not establish when the associations between organelles arise after target recognition or whether these organelle contacts arise at the synapse itself. We therefore followed the organelle network through individual killing events.

### A three-phase programme assembles and delivers the organelle ensemble

We next determined the order in which multiple organelles were trafficked to the immune synapse in individual CTLs. This is beyond the capability of conventional live-cell imaging as i) fluorophores are typically imaged sequentially, leading to colour drift artefacts, and ii) fluorophores have relatively broad emission spectra leading to spectral overlap, limiting the number of components which can be accurately imaged in the same cell. Recently we developed an eight-camera, multispectral acquisition module to acquire up to eight fluorophores simultaneously in live cells, compatible with any camera-based microscope [26]. We lever-aged this imaging architecture, alongside RLSU, to characterise changes to the organelle interactome volumetrically in CTLs during killing at uncompromised spatiotemporal resolution (Figure 3a).

**Figure 3:**
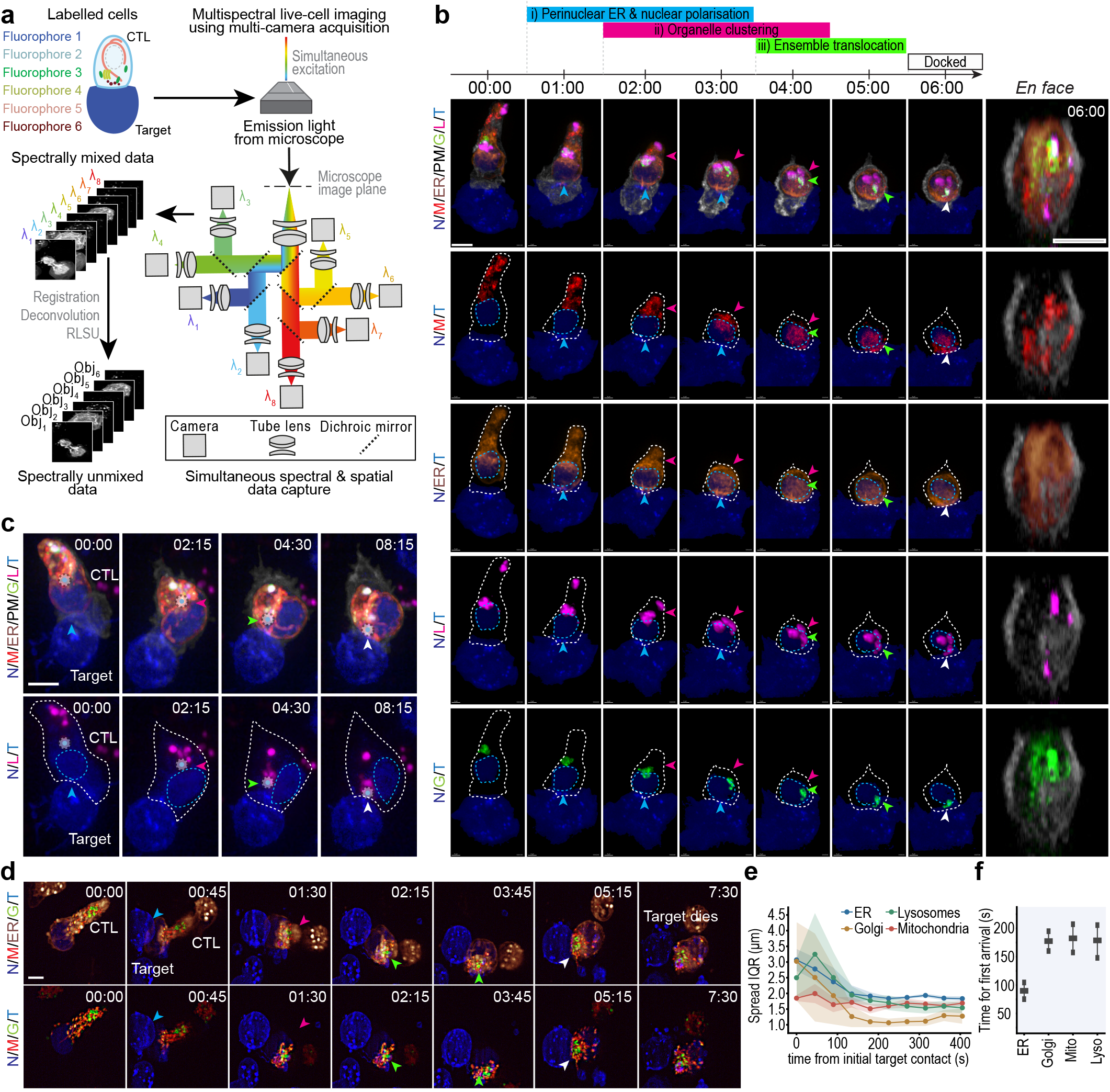
Organelles cluster distally around the centrosome and are delivered to the synapse as a multi-organelle ensemble. **a** Schematic of live-cell multispectral volumetric imaging of organelles in individual CTLs during killing events using a multi-camera imaging module followed by RLSU processing. CTLs with combinations of organelles labelled by polycistronic plasmids and/or live imaging dyes were excited with several lasers simultaneously, the resulting emission from up to eight fluorophores was imaged simultaneously using a tree-like arrangement of dichroic mirrors and eight synchronised cameras, as described in [26]. The resulting volumes were spatially registered, deconvolved and spectrally unmixed using RLSU. **b** Multispectral live imaging of a representative OT-I CTL recognising and killing an OVA-pulsed EL4 target, aligned to initial target recognition. Maximum intensity projections are shown with corresponding combinations of organelles as shown in row labels. Final column depicts *en face* views of the immune synapse formed after 6 minutes following target engagement. Timeline with corresponding events is shown: merge and single-channel views illustrating target recognition, early perinuclear ER/nuclear polarisation (cyan arrow), organelle clustering at the cell rear (magenta arrow), translocation around the nucleus (green arrow) and final synaptic docking (white arrow). Dashed grey line, CTL plasma membrane; dashed blue line, nucleus. **c** Centrosome-labelled killing sequence showing early perinuclear ER/nuclear polarisation (cyan arrow), organelle clustering around the centrosome at the cell rear (magenta arrow), translocation of the centrosome-organelle ensemble around the nucleus (green arrow) and final synaptic docking (white arrow). Bottom panel ER, mitochondria, Golgi and plasma membrane channels omitted. **d** Multispectral oblique-plane light-sheet imaging of the same three-phase programme over a complete killing event (target death is shown in the final frame). Cyan arrow, proximity between perinuclear ER and target immediately after recognition; magenta arrow, organelle compaction; green arrow, translocation of multi-organelle ensemble around nucleus; white arrow, multi-organelle ensemble reaches IS. Bottom panel ER channel omitted. **e** Spatial spread of ER, Golgi, lysosomes and mitochondria, quantified as interquartile range during killing events; note compaction from 120 s after target recognition. **f**, First-arrival time of each organelle at the synaptic plasma membrane (distance ≤ 500 nm). **e-f**, Mean and s.e.m. (*n* = 6 cells, *N* = 3 independent experiments). Maximum intensity projections (**b-d**), (*n* = 6 cells, *N* = 3 independent experiments). N, nucleus, blue; M, mitochondria, red; ER, orange; G, Golgi, green; L, lysosomes, magenta; T, target cell, blue; PM, plasma membrane, grey. IS, immune synapse.

Multispectral volumetric live-cell imaging resolved an ordered sequence of intracellular changes following target recognition (Figure 3b-d, Supplementary Video 4). Time-lapses were temporally aligned to the moment of initial target recognition, enabling us to reconstruct the sequence of intracellular events following TCR engagement across individual killing events relative to a common biological landmark.

Before target engagement, mitochondria, lysosomes, the Golgi, and ER occupied the rear of the migrating CTL, behind the nucleus, consistent with the migratory architecture observed in fixed cells. Target recognition initiated a rapid and characteristic reorganisation that could be delineated into three phases.

#### Phase 1: perinuclear ER and nuclear polarisation

Within approximately one minute of target recognition, the perinuclear ER extended towards the nascent synaptic interface, followed by polarisation of the nucleus towards the target (Figure 3b,c). Peripheral tubular ER, alongside other organelles, remained concentrated at the cell rear. The perinuclear ER was therefore the earliest organelle compartment to approach the target interface, with a mean earliest synaptic arrival time of approximately 100 s (Figure 3f). This places ER reorganisation alongside the recently described nuclear-polarisation step that precedes centrosome docking [11].

#### Phase 2: organelle assembly at the centrosome

Over the next several minutes, coincident with uropod retraction, the peripheral ER, Golgi, mitochondria and lysosomes trafficked towards the centrosome behind the nucleus. These organelles became increasingly clustered from approximately 120 s after target recognition (Figure 3e) and formed an ensemble around the centrosome (Figure 3c).

#### Phase 3: synaptic delivery of the centrosomal multi-organelle assembly

Following centrosomal clustering, the centrosome-associated organelle assembly translocated around the nucleus toward the immune synapse. Golgi, mitochondria and lysosomes arrived within a similar temporal window, consistent with coordinated delivery of the compacted multi-organelle architecture rather than independent sequential arrival (Figure 3b-c,f). The same sequence was observed by multispectral oblique-plane light sheet imaging, which allowed complete killing events to be captured up to and including target cell death (Figure 3d, Supplementary Video 5, Supplementary Video 6).

Thus, target recognition reorganises CTL organelles through a characteristic programme (COMET) in which an early ER/nuclear response is followed by centrosome-associated organelle clustering and subsequent synaptic delivery of the clustered multi-organelle ensemble with the centrosome. Together, these observations extend the established choreography of actin depletion, centrosome polarisation and granule recruitment [5, 6, 8, 9, 27] into a three-phase programme that encompasses the broader intracellular organelle network.

### The organelle interactome is remodelled before delivery to the synapse

We next asked when and where during polarisation organelle contacts formed. We therefore adapted our volumetric contact analysis pipeline to quantitate organelle contacts in time and space, in relation to the immune synapse. Object-based contact detection was applied to segmented unmixed multispectral live-cell volumes, and pairwise contact volumes were quantified through individual CTL-target encounters (Figure 4a-f). Values were normalised to the frame of initial target recognition (t_0_), so unity represents the interaction level at engagement rather than absence of interaction.

**Figure 4:**
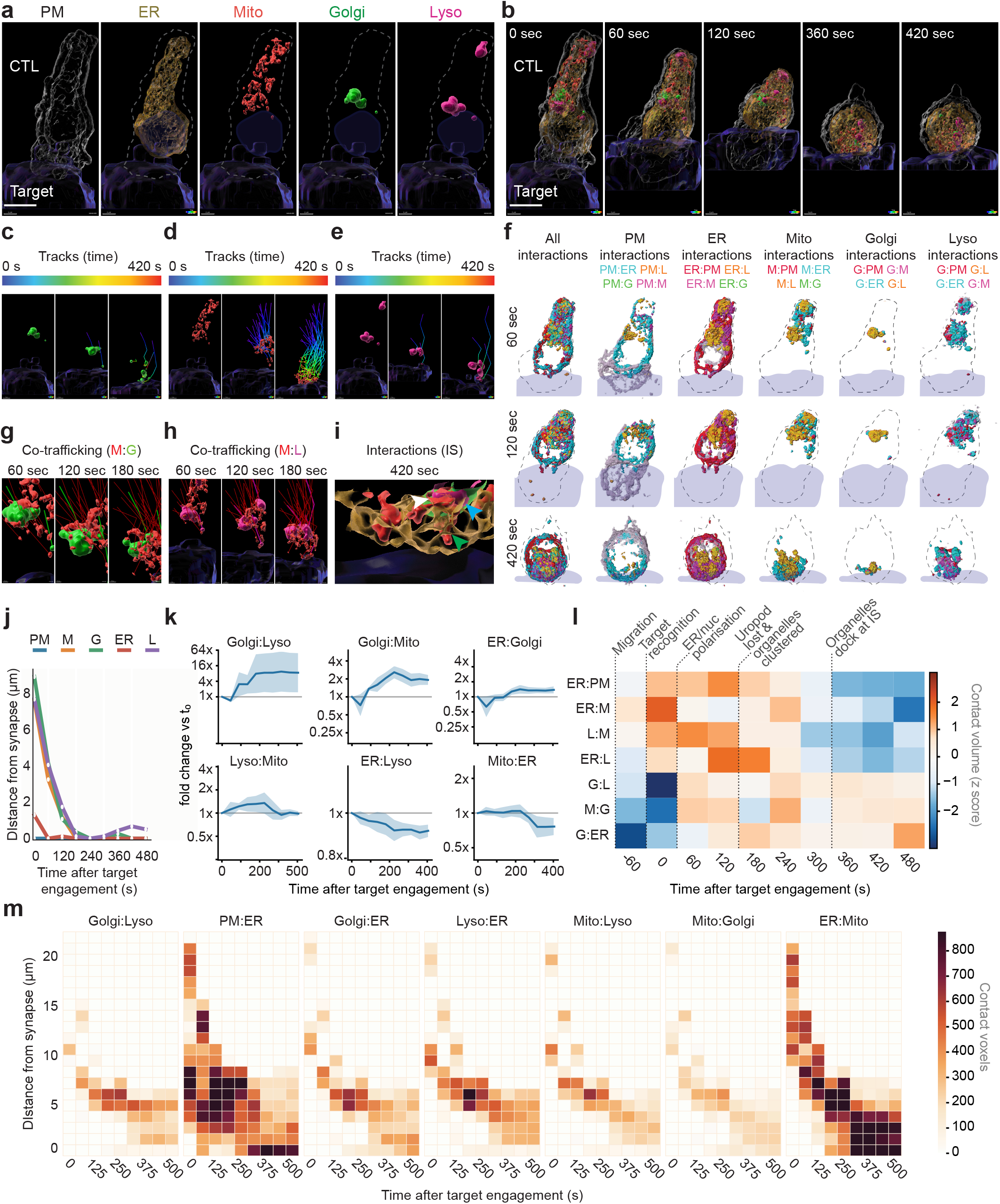
The CTL organelle interactome is rewired distally and delivered to the synapse. **a-m**, Multispectral live spinning-disk confocal imaging of organelle interactions in OT-I CTLs during killing. Timelapses referenced to initial target recognition (*t*_0_). **a, b**, Organelle surfaces and merged time series. Dashed line, CTL boundary. **c-e**, 3D tracks (0-420 s, colour-coded by time) of Golgi (**c**), mitochondria (**d**) and lysosomes (**e**). **f**, Contact maps at 60, 120 and 420 s, for all contacts (left) and split by participating organelle. Grey, underlying organelle mask. **g-h**, Overlaid tracks showing co-trafficking of mitochondria with Golgi (**g**) and lysosomes (**h**). **i**, Organelle contacts (segmentations) at the immune synapse at 420 s. **j**, Organelle distance from the synapse over time (representative cell). **k**, Pairwise contact volume as fold change relative to *t*_0_(1 = level at target recognition). (*n* = 6 cells, *N* = 3 independent experiments; mean and 95% confidence interval). **l**, Within-pair z-scored temporal profiles with morphological landmarks annotated; z-scores do not rank pairs by absolute abundance. **m**, Contact voxels (kymograph) by distance from the synapse and time for seven organelle pairs (representative cell). PM, plasma membrane; ER, endoplasmic reticulum; M, mitochondria; G, Golgi; L, lysosomes; IS, immune synapse. Scale bars, 5 µm.

Three-dimensional tracking confirmed directional movement of the Golgi, mitochondria and lysosomes towards the synapse (Figure 4c-e). Subsets of mitochondrial tracks over-lapped spatially with Golgi or lysosome trajectories over multiple frames, consistent with partial co-trafficking of these compartments during reorganisation (Figure 4g-h). By the end of the analysed interval, extensive ER-, Golgi-, lysosome- and mitochondria-associated contacts were concentrated at the immune synapse (Figure 4i,j).

To determine whether the interactome is remodelled during killing, we asked: i) how much each contact changes, ii) when it changes relative to the three-phase programme, and iii) where in the cell the changes occur. Expressing organelle contact volume as fold change relative to t_0_ revealed Golgi-lysosome contacts rose steeply from approximately 90 s, exceeding four-fold within the first three minutes of target engagement and plateauing at approximately eight-fold thereafter (Figure 4k). Golgi-mitochondria contacts followed a similar trajectory of smaller amplitude, peaking above two-fold at approximately 180 s before settling near two-fold, and ER-Golgi contacts stabilised approximately 1.3-fold above the engagement level. Lysosome-mitochondria contacts rose moderately, peaking at approximately 1.35-fold between 135 and 225 s, and returned to their engagement level by 315 s. ER-lysosome contacts declined progressively to approximately 0.75-fold before stabilising. ER-mitochondria contacts were maintained at their engagement level across the first 225 s, and then fell sharply between 225 and 315 s to approximately 0.8-fold, where they remained. Importantly, a uniform compaction of the cytoplasm would increase all contacts in proportion, yet ER-lysosome contacts fall and lysosome-mitochondria contacts return to baseline.

Normalising each organelle pair across time further revealed successive contact configurations aligned with the three-phase morphological programme (Figure 4l). ER-associated contacts were relatively enriched during early polarisation and centrosomal assembly, whereas Golgi-centred relationships became more prominent as the organelle architecture approached and reached the synapse.

Finally, we decomposed organelle contacts jointly by time after target recognition and distance from the synapse (Figure 4m). In representative cells, across the major pairs analysed, enrichment first appeared at distal intracellular positions and subsequently traced a trajectory towards the 0–2 µm region adjacent to the synapse. These data are consistent with contact-rich organelle arrangements being established during the intracellular reorganisation and then redistributed towards the synapse, rather than being generated exclusively after synaptic arrival.

### ER calcium-store depletion promotes centrosomal organelle assembly but not synaptic delivery

The early behaviour of the perinuclear ER and the changing distribution of ER-associated contacts prompted us to examine ER-plasma membrane contact sites during CTL killing. To visualise these ER-PM contact sites we expressed MAPPER [28], a genetically encoded reporter that accumulates at ER-PM junctions. MAPPER is anchored in the ER using the transmembrane domain of STIM1 with a carboxy-terminal polybasic motif that reversibly binds to negatively charged lipids in the PM if it is within 25 nm from the ER [28]. In migrating CTLs, MAPPER-positive structures were concentrated pre-dominantly toward the cell rear. Following target recognition, MAPPER signal appeared rapidly near the nascent synaptic interface within areas of actin depletion, while additional MAPPER-positive clusters accumulated at the cell rear and subsequently redistributed towards the synapse over several minutes (Figure 5c, d, Supplementary Video 7). These observations reveal rapid remodelling of ER-PM contact sites during the same interval in which the wider organelle network is reorganised. However, as MAPPER reports close ER-PM apposition rather than the molecular composition or activity of individual junctions, we do not infer that all MAPPER-positive structures are functionally equivalent.

**Figure 5:**
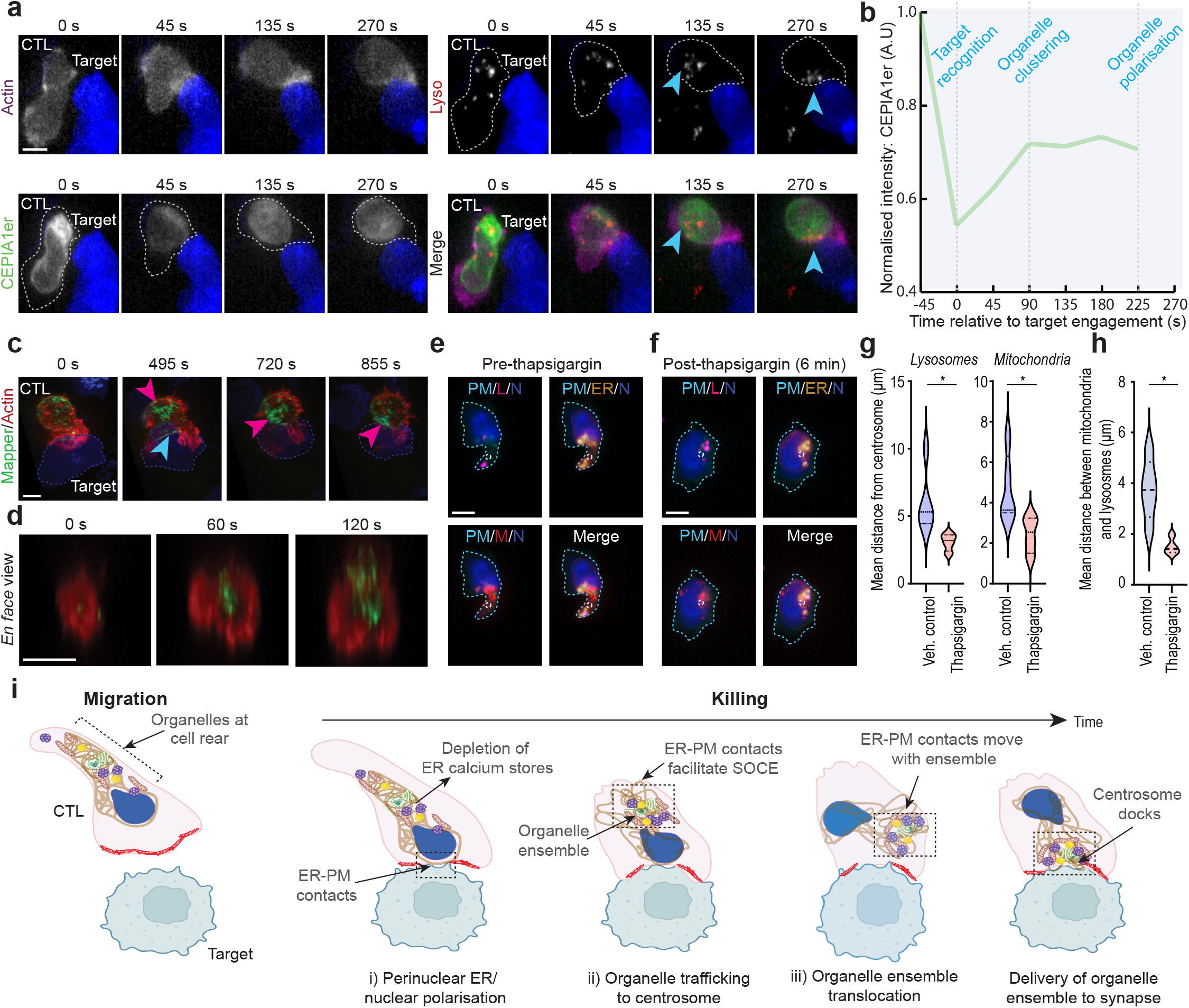
ER-PM remodelling and ER calcium store depletion precede centrosomal organelle assembly. **a** Actin, G-CEPIA1er and lysosomal channels, with merged views, during CTL target engagement. Dashed grey line depicts CTL cell boundary. Representative cell (*n* = 4 cells, *N* = 3 independent experiments). **b** Normalised G-CEPIA1er fluorescence relative to the pre-recognition signal, shown alongside morphological landmarks. The trace reports fluorescence rather than a calibrated fraction of stored calcium. Time = 0 represents the time-point of initial target recognition. Trace from a representative cell (*n* = 4 cells, *N* = 3 independent experiments). **c** MAPPER and actin during target engagement. Cyan and magenta arrows indicate reporter accumulation near the synapse and at the cell rear, respectively. Representative cell (*n* = 5 cells, *N* = 3 independent experiments). **d** *En face* view of actin (red) and MAPPER (green) over time at the IS. **e, f** A representative CTL imaged before (e) and 6 min after (f) addition of 0.6 *µ*M thapsigargin. Dashed white circle indicates the centrosome. **g** Mean distance of lysosomes (left) and mitochondria (right) from the centrosome in independent CTLs imaged 6 min after addition of vehicle (DMSO) or 0.6 *µ*M thapsigargin; each point is one cell (*n* = 5 cells per condition, *N* = 3 independent experiments). P = 0.0456 and 0.0407, respectively. **h** Mean distance between mitochondria and lysosomes in the same cells (*n* = 5 cells per condition, *N* = 3 independent experiments). P value = 0.0117. Statistics: two-sided Welch’s unpaired t-test. PM, plasma membrane; N, nucleus; M, mitochondria; L, lysosomes; ER, endoplasmic reticulum. **i** Schematic depicting intracellular events following target recognition.

ER-PM junctions are also where store-operated calcium entry (SOCE) takes place: depletion of ER Ca^2+^ drives STIM1 to accumulate at these contacts, where it binds and activates plasma membrane ORAI1 channels [20–23]. The remodelling of ER-PM contacts we observed therefore raised the question of how ER store dynamics are timed relative to the organelle programme. We therefore imaged the ER-luminal Ca^2+^ reporter G-CEPIA1er [29] together with actin and lysosomes during target engagement (Figure 5a, Supplementary Video 8). Target recognition was followed by synaptic actin depletion, perinuclear ER polarisation and an approximately 50% decrease in G-CEPIA1er fluorescence (Figure 5a,b). The decrease in ER calcium signal preceded lysosomal clustering and polarisation, and the ER signal subsequently recovered towards approximately 70% of its pre-engagement level which coincided with lysosomal clustering (Figure 5a,b). This temporal ordering placed ER calcium-store depletion before centrosomal organelle clustering and synaptic delivery.

We next asked whether ER Ca^2+^ store depletion is *sufficient* to promote the early architectural response in the absence of target cells. Migrating CTLs were acutely treated with thapsigargin, a potent inhibitor of sarco/endoplasmic reticulum Ca^2+^ ATPases, that depletes ER Ca^2+^ thereby increasing cytosolic calcium [30]. Within 6 min, thapsigargin induced trafficking of organelles towards the centrosome (Figure 5e,f). Specifically, mitochondria and lysosomes moved significantly closer to the centrosome, and each other, in thapsigargin-treated cells compared with vehicle-treated cells (Figure 5g,h). However, thapsigargin did not reproduce the directional relocation of the assembled architecture towards the cell front observed during productive target encounters.

These data separate two organisational steps in the cytotoxic response. ER calcium-store depletion is sufficient to promote substantial centrosomal organelle recruitment, but an additional spatially instructive process is required for directional delivery of the assembled architecture to the immune synapse.

## Discussion

Our findings identify COMET and place CTL secretory granule polarisation within a broader systems-level reorganisation of the organelle network [1–3]. Target recognition initiates early perinuclear ER and nuclear polarisation, multiple organelles then converge around the centrosome on the distal side of the nucleus. The resulting organelle ensemble moves with the centrosome around the nucleus to the synapse. The immune synapse is therefore not simply a destination to which independently trafficked compartments arrive but rather the delivery point of an assembled and repositioned multi-organelle unit.

MTOC and Golgi reorientation was recognised early as a hallmark of CTL-target engagement [4]. Subsequent work established centrosome docking as a direct route for lytic granule delivery [2], showing that granule recruitment to the centrosome can be regulated independently of centrosome polarisation [5] and separating centrosome translocation from the final docking step [6, 31]. Four-dimensional imaging then ordered actin depletion, centrosome movement and granule secretion [8, 9, 27], while nuclear polarisation was recently placed before centrosomal docking [11]. Our data now show that the same response encompasses the ER, Golgi, mitochondria and lysosomes as a coordinated organelle ensemble. In this view, the centrosome is not only a guide for secretory lysosomes but also an organising hub around which a broader intracellular architecture is dynamically assembled. The organelle relationships within that architecture are dynamic. Electron tomography demonstrates abundant membrane contact sites among synapse-polarised organelles, while live imaging shows that contacts change throughout the preceding intracellular reorganisation. Importantly, these relationships are not confined to the synapse: contacts emerge at distal intracellular positions and are subsequently redistributed towards the synapse with the organelle ensemble. We demonstrate selective remodelling of organelle relationships during the transition from migration to killing. As distinct organelle contacts predominate at successive stages of the killing process, it will be important to determine the functional role of these membrane contact sites in sustaining cytotoxicity.

Our results also suggest that assembly of the organelle ensemble and synaptic delivery are separable stages. Thapsigargin-induced ER store depletion [30] is sufficient to recruit mitochondria and lysosomes towards the centrosome, but does not recreate the directional movement of the assembled architecture towards the cell front. This is consistent with a model in which an early signal coupled to ER store depletion contributes to organelle assembly, whereas spatially instructive signals and cytoskeletal mechanisms direct the assembled state towards the target.

The early ER response connects the organelle programme to established T-cell Ca^2+^ biology. STIM1 and ORAI1 couple ER store depletion to Ca^2+^ entry at ER-plasma membrane junctions [20–23], and can accumulate at the immune synapse and in distal caps during T-cell activation [12–14], while mitochondria positioned near the synapse could help shape local Ca^2+^ microdomains [14, 15]. Here, MAPPER-labelled ER-PM contact sites [28] and ER Ca^2+^ levels [29] change during the same early period in which the perinuclear ER moves towards the target and the broader organelle network begins to cluster. The data therefore place ER-PM remodelling within a cell-wide architectural response. Determining which ER-PM contact sites contain STIM/ORAI complexes, how their molecular composition changes over time, and whether individual contact sites are required for subsequent organelle assembly will require direct molecular perturbation. Beyond STIM1 and ORAI1, ER-PM contacts also host lipid-transfer proteins such as E-Syt1 and Nir2, which replenish plasma membrane PI(4,5)P_2_ after receptor-triggered PLC activity [28]; whether such machinery tunes the DAG and phosphoinositide landscape of the synapse, thereby linking ER-PM remodelling to centrosome polarisation, is an important question for future work.

Additionally, the organelle ensemble model generates testable functional predictions. CTLs are serial killers and can replenish cytotoxic machinery while moving between successive targets [32]; termination of TCR signalling and target disengagement are themselves actively regulated at the synapse [33]. If a multi-organelle architecture remains partially assembled after one killing event, repolarisation towards a second target could occur more rapidly than *de novo* assembly. Conversely, as relocating an assembled organelle ensemble around the nucleus imposes both a delay and a mechanical cost, the programme could act as a threshold that weak or transient engagements fail to cross, restricting secretion to productive contacts and limiting bystander killing.

Together, our findings establish a framework for understanding how dynamic organelle organisation underpins specialised cellular functions. Combining high-dimensional live imaging with targeted perturbations should reveal how these collective states are established, maintained and dismantled, and whether analogous programmes govern other cellular behaviours. More broadly, this work positions the organelle interactome as a dynamic and fundamental level of organisation through which cells execute complex functions.

## Methods

### Mice

C57BL/6 (B6)-OT-I Rag2^−/−^ [B6.129S6-Rag2^tm1Fwa^ Tg (TcraTcrb)1100Mjb] mice >6 weeks old were bred and housed in the University of Cambridge Facility. This research has been regulated under the Animals (Scientific Procedures) Act 1986 Amendment Regulations 2012 following ethical review by the University of Cambridge Animal Welfare and Ethical Review Body (AWERB).

### Cell culture

Naïve OT-I splenocytes were stimulated with 10 nM ovalbumin (OVA) peptide, SIINFEKL, for 3 days in CTL medium: RPMI 1640 medium with 10% fetal bovine serum (FBS), 50 µM *β*-mercaptoethanol, recombinant murine interleukin-2 (10 U/ml), 2 mM L-glutamine, 1 mM sodium pyruvate, penicillin (50 U/ml), and streptomycin (50 µg/ml). Cells were pelleted and resuspended in fresh medium daily from 3 days after stimulation, expanded in CTL medium, and used at 6 to 9 days after stimulation. EL4 target cells stably expressing the mTagBFP2-Farnesyl-5 (mTagBFP, monomeric blue fluorescent protein; EL4-blue) were cultured in Dulbecco’s Modified Eagle Medium (DMEM) with 10% FBS. Four to 24 hours before live-cell imaging or EM sample preparation, 5 to 10 × 10^6^ CTLs were nucleofected with 2.5 µg of DNA per plasmid, using the Amaxa nucleofector (programme DN-100) with Lonza P3 Primary Cell 4D-Nucleofector Kit. Plasmids and dyes used in this study are listed in Supplementary Table 1. In some instances, live-cell dyes, such as SPY 555 FastAct_X and SiR lysosomes (Spirochrome) were used. Lyophilised live-cell dyes were reconstituted in 50 µL dimethyl sulfoxide to prepare a 1000× stock solution according to manufacturer’s instructions. The stock solution was stored at −20 °C. On the day of the experiment, the probe was diluted to a 1× staining solution in full growth medium and vortexed briefly. OT-I were pelleted and resuspended in staining solution for one hour at 37 °C. The cells were subsequently pelleted and washed twice with PBS, prior to resuspension in warmed imaging buffer [RPMI without phenol red, 10% FBS, 25 mM Hepes, 2 mM L-glutamine, penicillin (50 U/ml), and streptomycin (50 µg/ml)].

### Live-cell imaging sample preparation

Glass-bottom imaging dishes (World Precision Instruments, FD35 or MatTek, P35G-1.5-14-C) were coated with intercellular adhesion molecule-I (ICAM-1; 0.5 µg/ml) in phosphate-buffered saline (PBS) at 4 °C overnight and gently washed with PBS just before use. For killing experiments, target cells (EL4-blue) were pulsed with 1 µM OVA for 1 hour before washing, resuspended at 6.5 × 10^5^ cells/ml in serum-free DMEM, plated in the glass area of dishes (0.3 mL per dish), and allowed to attach to the glass at 37 °C. After 5 min, 1.5 mL of warmed imaging buffer [RPMI without phenol red, 10% FBS, 25 mM Hepes, 2 mM L-glutamine, penicillin (50 U/ml), and streptomycin (50 µg/ml)] was used to disperse un-attached target cells from the glass area. The target cell dish was mounted on the microscope stage in an environmental chamber at 37 °C and 5% CO_2_ (Okolab) for imaging before CTLs labelled with live-cell imaging dyes and/or expressing fluorescent-tagged proteins in imaging buffer were added dropwise to the dish of target cells. For studies of migrating CTLs, OT-I cells were resuspended in serum-free RPMI without phenol red and added dropwise to coated glass-bottomed dishes at 37 °C. After 10 minutes, imaging buffer was added to the cells and the cell dish was mounted on the microscope stage in an environmental chamber at 37 °C and 5% CO_2_ (Okolab) for one hour prior to imaging.

### Fixed-cell imaging sample preparation for multiplexed Airyscan imaging

CTLs, expressing fluorescent proteins and/or labelled with live-cell imaging dyes, and SIINFEKL pulsed-target cells (EL4-blue) were mixed in serum-free medium (DMEM) and allowed to adhere to an ICAM-1-coated (0.5 µg/ml) glass eight-well chamber (Ibidi, 80841) for 5 min at 37 °C before adding medium with serum for 10 min prior to fixation. Migrating CTLs were pelleted in serum-free medium and added drop-wise to an ICAM-1-coated glass eight-well chamber and allowed to adhere for 5 min at 37 °C before adding medium with serum for 30 min prior to fixation. Cells were then fixed with pre-warmed 4% paraformaldehyde in PBS for 10 min and washed with PBS prior to immediate Airyscan imaging.

### Electron microscopy and tomography

CTL-target conjugates were formed at 37 °C, after which they were fixed in pre-warmed fixative containing 2% PFA (Electron Microscopy Sciences) and 1.5% glutaraldehyde (Agar Scientific) for 20 min at RT, then 30 min at 4 °C. The samples were maintained at 4 °C for all subsequent steps until dehydration to minimise diffusion of the cytochemical reaction product. Fixed samples were washed in 0.1 M sodium cacodylate, pH 7.4, rinsed in Tris buffer (0.5 M Tris HCl, pH 7.6), and processed for diaminobenzidine (DAB) cytochemistry by incubating for 15 min in DAB reaction mixture (DAB (Sigma) (0.75 mgmL^−1^), 0.21% (v/v) hydrogen peroxide in Tris buffer) in the dark. Samples were washed in Tris buffer followed by 0.1 M sodium cacodylate, pH 7.4, then post-fixed in 0.1% osmium tetroxide (TAAB) for 1 h followed by 0.5% aqueous uranyl acetate (TAAB) overnight, washing with deionised H_2_O in between. Post-fixed samples were dehydrated in increasing concentrations of ethanol (50-100%) and embedded in EPON resin (Agar Scientific).

For TEM tomography, 250-300 nm serial sections across the immune synapse were collected on formvar-coated slot grids, and the top and bottom of the sections decorated with 15 nm fiducial gold (British Biocell International). Tilt series images of cells of interest were collected manually for each serial section at 2° intervals from −60° to 60° using an FEI Tecnai Spirit TEM (FEI, Eindhoven) with a Gatan camera. Tilt series images were converted into .mrc series files using Cygwin, and tomograms were generated and serial tomograms joined using the IMOD and 3dmod packages (Boulder Laboratory for 3-D EM of Cells, University of Colorado, USA). Models of the joined tomograms were generated using 3dmod (Boulder). ImageJ was used for additional image processing and movie generation.

### Microscopy

#### Excitation-multiplexed Airyscan microscopy

Fixed samples were imaged on a Zeiss LSM 980 with an Airyscan 2 detector, using a 63×/1.40 NA Plan-Apochromat oil-immersion objective. Excitation-multiplexed volumes were acquired by recording a complete *z*-stack at each of six excitation wavelengths in turn (405, 445, 488, 514, 590 and 639 nm). The emission configuration was determined in ‘Smart Setup’ for the fluorophores used in the sample. Due to the overlapping spectra of the fluorophores used in the sample, each volume contained mixed signal from every fluorophore excited at each wavelength. Volumes were acquired in super-resolution mode with a lateral pixel size of 30 nm and a *z*-step of 130 nm, meeting Nyquist sampling for Airyscan reconstruction. Lateral resolution, measured with 100 nm TetraSpeck beads (T7279, Invitrogen), was 120-200 nm FWHM, compared with 250-400 nm for confocal acquisition on the same system.

#### Andor spinning disk system: SDCM

This system comprised a Leica DMi8 stand, a CSU-X1 spinning disk unit (Yokogawa), a Borealis illumination unit (Andor) with 405, 488, 561 and 640 nm lasers, and a TuCam dual-port adapter (Andor). Objectives were a UPLSAPO 100×/1.40 NA oil (Olympus), a 60×/1.30 NA oil (Olympus), an HC PL APO 100×/1.40 NA oil CS2 (Leica), an HC PL APO 63×/1.40 NA oil CS2 (Leica) and an HC PL APO 63×/1.30 NA glycerol CORR CS2 (Leica). One TuCam port carried an iXon Ultra 888 EMCCD camera (Andor). The other carried a custom eight-camera multispectral acquisition module [26]. A fully reflective flip mirror sent the emission to one port or the other, so the same microscope could be used in two modes.

In conventional mode, the flip mirror was withdrawn. Channels were acquired sequentially on the iXon Ultra 888 through 452/45, 525/50, 600/50 or 685/40 nm emission filters, under the control of Fusion software (Andor). This was used for experiments with up to four labels, including MAP-PER/actin and G-CEPIA1er/actin/lysosome imaging.

In multispectral mode, the flip mirror was inserted to send the emission to the eight-camera module, and all fluorophores were excited and recorded simultaneously through a quad-band dichroic. The module split the emission into eight spectral bands with a tree of dichroic mirrors, each band recorded by its own camera (xiX, Ximea). The cameras were connected through an xSWITCH (Ximea), hardware-triggered through their general-purpose inputs (GPI), and streamed over PCIe to a dedicated acquisition PC running Micro-Manager.

#### Excitation-multiplexed spinning disk system: SDCM-M

This system comprised a CrestOptics X-Light V3 spinning disk unit on a Nikon Ti2 stand, with a fast piezo *z*-stage (ASI), a Prime 95B back-illuminated sCMOS camera (Photometrics) and a 100×/1.40 NA Plan Apochromat oil-immersion objective. Illumination came from a multiline solid-state laser source (LDI-7, 89 North) with lines at 405, 445, 470, 520, 528, 555 and 640 nm. For excitation-multiplexed acquisition, the excitation wavelength was cycled at each volume. The system was controlled by Micro-Manager.

#### Multispectral oblique-plane microscopy: OPM

Light-sheet timelapses were acquired on a custom multispectral oblique-plane microscope (OPM), described in detail previously [26]. Fluorescence was collected by a 100×/1.35 NA silicon-immersion primary objective (Nikon). A remote-focusing relay then imaged a plane tilted at 30° to the optical axis onto the custom-built eight-camera multi-spectral module. The light sheet and image plane were scanned through the sample with a galvanometric mirror. Samples were excited with 405, 488, 561 and 638 nm lasers. The microscope was controlled by Micro-Manager and hardware-synchronised with a National Instruments DAQ card (PCIe-6321).

#### Live imaging conditions

All live imaging was performed at 37 °C and 5% CO_2_ in an environmental chamber (Okolab), in imaging buffer. For killing movies, timelapses were started before CTLs contacted targets. Time zero was defined as the first frame in which the CTL contacted its target. Microscopes used for each experiment are outlined in Supplementary Table 1.

### Image processing

All custom processing was implemented in Python 3.

#### Channel registration

Data from the eight-camera module were registered to remove the small offsets between cameras. Reference images of back-illuminated lens tissue were acquired on all cameras for each imaging session. Transforms mapping each channel onto channel 1 were estimated from these images by DFT-based phase correlation `imreg_dft`), saved, and applied to the experimental data.

#### Deskewing

OPM stacks were deskewed with `lsfm_tools` using linear interpolation after registration.

#### Deconvolution

Live multispectral spinning disk data were deconvolved with the Richardson-Lucy algorithm (4 iterations) before unmixing. Each channel was deconvolved with a theoretical point spread function computed for its emission wavelength, objective NA, refractive index and voxel size. Airyscan data were reconstructed in ZEN Blue (Zeiss) by Airyscan joint deconvolution (jDCV), which combines pixel reassignment with accelerated Richardson-Lucy deconvolution. jDCV used the Standard sample profile, 5 iterations and a quality threshold applied identically to all channels.

#### Spectral unmixing

Spectrally mixed data were unmixed by Richardson-Lucy Spectral Unmixing (RLSU) [26] with 1000 iterations. RLSU solves for non-negative fluorophore abundances under a Poisson noise model, given a mixing matrix **M** whose element *M*_*i j*_is the fractional contribution of fluorophore *j* to channel *i* . For emission-multiplexed data, each channel is one camera’s emission band. *M*_*i j*_was calculated from the emission spectrum of fluorophore *j* and the transmission of the dichroics and filters routing light to camera *i* . For excitation-multiplexed data, each channel is one excitation wavelength recorded through a fixed emission path. *M*_*i j*_was calculated from the excitation efficiency of fluorophore *j* at wavelength *i*, weighted by the fraction of its emission passing the detection filters. Reference spectra were taken from FPbase and refined using single-label controls and pixels belonging unambiguously to a single labelled structure.

#### Processing pipelines

Excitation-multiplexed Airyscan: jDCV, then RLSU. Multi-camera spinning disk: registration, then RL deconvolution, then RLSU. Multi-camera OPM: deskewing and registration, then RLSU. Excitation-multiplexed spinning disk: RL deconvolution, then RLSU. Conventional spinning disk: no deconvolution; no unmixing.

#### Validation of excitation-multiplexed unmixing

Unmixing for the six-fluorophore Airyscan labelling scheme was validated in simulation (Supplementary Figure S1). Six spatially distinct ground-truth objects were each assigned one fluorophore and forward-projected through the calculated mixing matrix. Poisson noise was added to reproduce photon-limited acquisition. The result was then unmixed with RLSU. Accuracy was quantified as the root-mean-square error (RMSE) between unmixed and ground-truth objects. The mixing matrix had a condition number of 3.16, and the maximum RMSE across the six channels was 1.13.

### Quantification of organelle position, clustering and synaptic arrival

Organelles were segmented in Imaris (v11.0, Oxford Instruments). Mitochondria, Golgi and lysosomes were segmented with the Surfaces function. The PM and ER were segmented with the Machine Learning Segmentation function, using a pixel classifier trained on manually annotated organelle and background voxels. Merged punctate objects (mitochondria, lysosomes) were separated by watershed splitting. Masks were exported as 3D binary volumes and analysed in Python 3 using a custom analysis script. All live measurements used an absolute time axis, with *t* = 0 defined as the first frame of contact between the CTL and its target.

The synaptic plasma membrane was defined at each time-point as the voxels shared by the CTL and target-cell PM masks.

#### Organelle distribution along the rear-to-front axis

For each cell, a rear-to-front axis was defined and cells were aligned along this axis. For migrating CTLs, it ran from the rear of the uropod to the leading edge. For CTL-target conjugates, it ran from the cell rear to the centroid of the synapse. Every voxel of each organelle mask was projected onto this axis. Positions were normalised to cell length, from 0 at the rear to 1 at the front, so that cells of different sizes could be compared. The normalised axis was divided into 100 equal bins. In each bin, the summed fluorescence of each channel was expressed as a fraction of that channel’s total in the cell, so each profile sums to one. Profiles were then averaged across cells for each condition. To quantify organelle volume ahead of the nucleus, we calculated the fraction of each organelle’s segmented volume that projected beyond the front-most point of the nuclear mask.

#### Organelle clustering

Organelle spread was measured at each timepoint as the interquartile range (IQR) of the Euclidean distances of all voxels of a given organelle mask from that mask’s centroid. A decrease in IQR indicates compaction of that organelle. Values were averaged across cells at each timepoint (mean ± s.e.m.).

#### Synaptic arrival time

For each organelle at each time-point, we measured the segmented volume lying within 500 nm of the synaptic plasma membrane, using a Euclidean distance transform of the synapse mask that accounts for anisotropic voxel size. First-arrival time was defined as the first timepoint at which this volume exceeded 0.2 µm^3^. The threshold excluded isolated segmentation fragments. Arrival times are reported as mean ± s.e.m. across cells.

#### Organelle distance from the synapse

At each timepoint, the distance of every voxel of an organelle mask to the nearest synaptic voxel was computed. These were summarised per organelle as the median distance.

### Quantification of organelle contacts

Organelle contacts were quantified with an object-based approach using a custom-written script, implemented in Python 3 `numpy`, `scipy`, `scikit-image`). It was applied to 3D segmentation masks of the plasma membrane (PM), ER, Golgi, mitochondria and lysosomes, generated as described above. Because the distances involved are below the resolution of light microscopy, we refer to the measured quantities as organelle contacts. Only appositions confirmed by electron tomography are described as membrane contact sites.

#### Mask preparation

For each cell, the masks were restricted to the CTL, using the CTL PM mask filled in 3D; target-cell signal was excluded. Merged punctate objects (mitochondria, lysosomes) were separated by distance-transform watershed splitting before analysis. For each pair of organelles *A* and *B*, a voxel of *A* was classed as in contact with *B* if it lay within a distance *d* of the mask of *B* . Distances were computed from Euclidean distance transforms in physical units, accounting for anisotropic voxel size. *d* was set to 120 nm for Airyscan data and 250 nm for spinning disk data, approximately the lateral FWHM of each system’s point spread function. For each pair, the contact (interfacial) volume was defined as

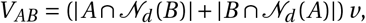

where *N*_*d*_(*X*) is the set of voxels within *d* of mask *X* and *v* is the voxel volume.

For fixed migrating CTLs and CTL-target conjugates, contacts were expressed as directed fractions. The fraction for *A* with *B*, *f* _*A*→*B*_= |*A* ∩ *N*_*d*_(*B*)|/|*A*|, is the percentage of the reference organelle *A*’s volume in contact with partner *B* . The matrix is therefore asymmetric. Values were averaged across cells for each condition. Colour-coded contact maps show voxels in contact, coloured by the participating organelle.

For live killing timelapses *V*_*AB*_was computed at every timepoint for the ten pairs among PM, ER, Golgi, mitochondria and lysosomes. All timelapses were analysed on an absolute time axis, with *t* = 0 at initial target recognition. To allow comparison across cells with different organelle loads, each pair’s trajectory was expressed as fold change relative to its value at *t* = 0, *V*_*AB*_(*t*)/*V*_*AB*_(0); a value of 1 therefore denotes the contact level at recognition, not the absence of contact. To compare the timing of changes between pairs independently of their amplitude, each pair’s trajectory was also z-scored across time within that pair. These values show when each pair is relatively enriched but do not rank pairs by absolute contact volume.

The immune synapse was defined from the overlap of the CTL and target PM masks, as described above. For every contact voxel, the Euclidean distance to the nearest synaptic voxel was computed. Contact voxels were then binned by distance from the synapse (bins of 1 µm) and by time, giving a distance × time map of contact volume for each pair. Organelle distance from the synapse was calculated in the same way from all voxels of each organelle mask.

### ER Ca^2+^ measurements

ER luminal Ca^2^+ was imaged in CTLs expressing G-CEPIA1er [29] during target engagement. At each timepoint, a whole-cell mask of the CTL was generated by Otsu thresholding of the actin channel, excluding the target cell. The integrated G-CEPIA1er intensity within the mask was measured over the whole volume. Background, measured in a cell-free region, was subtracted from each voxel before integration. Intensities were normalised to the mean of the pre-engagement frames, giving *F* /*F*_0_. No photobleaching correction was applied. Because G-CEPIA1er was not calibrated, *F* /*F*_0_reports relative changes in luminal Ca^2+^, not absolute store content. Morphological landmarks were scored from the actin and lysosome channels of the same timelapse and plotted alongside the trace. These were target recognition, organelle clustering and organelle polarisation to the synapse.

### Acute ER Ca^2+^ store depletion

Migrating CTLs transiently expressing PACT-TagBFP (centrosome), NLS-TagBFP (nucleus), Lyn-Cerulean (plasma membrane), MTS-mAzamiGreen (mitochondria) and mCherry-KDEL (ER) were co-stained with SiR Lysosome (Spirochrome) and plated on ICAM-1-coated dishes. Thapsigargin (0.6 µM final; T7459 Invitrogen) or vehicle (DMSO, 0.06% final) was added as a 10× solution in pre-warmed imaging buffer, gently and away from the imaging field. To illustrate the response, individual cells were imaged immediately before and 6 min after addition (Figure 5e,f). For quantification, independent thapsigargin- and DMSO-treated cells were imaged 5-10 min after addition (n = 5 cells per condition, pooled from N = 3 independent experiments). Mitochondria, lysosomes and the centrosome (PACT-TagBFP) were segmented in Imaris (v11.0). For each mitochondrion and lysosome, the Euclidean distance from its centroid to the centrosome was measured, and the distance between mitochondria and lysosomes was measured as the mean nearest-neighbour centroid distance. Distances were averaged per cell, each cell was treated as the statistical unit, and groups were compared using a two-sided Welch’s unpaired t-test.

### Statistics and reproducibility

Biological and technical sample sizes are reported in the figure legends where available. Unless otherwise indicated, n denotes individual cells and N denotes independent experiments. Analyses were performed in Python 3 (SciPy) or GraphPad Prism.

## Data and code availability

Data and code supporting the findings of this study are available from A.K. on reasonable request.

## Acknowledgements

This work was supported by a Wellcome Discovery Award (306545/Z/23/Z) provided to G.M.G. A.K. thanks E. Derivery and J. Manton for access to imaging infrastructure at the MRC Laboratory of Molecular Biology, the CIMR microscopy facility for access to the Zeiss LSM 980 Airyscan used in this work, the CIMR computing facility for their assistance, the Griffiths lab for interesting discussions related to this work, as well as Fitzwilliam College Cambridge and the Isaac Newton Trust for providing funding/support (Early Career Research Fellowship). G.M.G. and A.K. thank Pablo Carbajal Martin for tracing electron tomograms provided by J.C.S.

## Author contributions

A.K. and G.M.G. conceptualised the study. A.K. developed and implemented the multispectral imaging and computational analysis approaches, performed the fluorescence microscopy experiments, analysed the data and prepared the figures. J.C.S. performed the electron microscopy and electron tomography experiments and associated ultrastructural analysis. G.M.G. supervised/funded the research and provided scientific guidance. A.K. wrote the original manuscript with contributions from G.M.G. A.K. and G.M.G reviewed and approved the final manuscript.

