## Supplemental Files for "Systems-level imaging reveals assembly and synaptic delivery of a multi-organelle ensemble during T cell killing"

### Supplemental figures

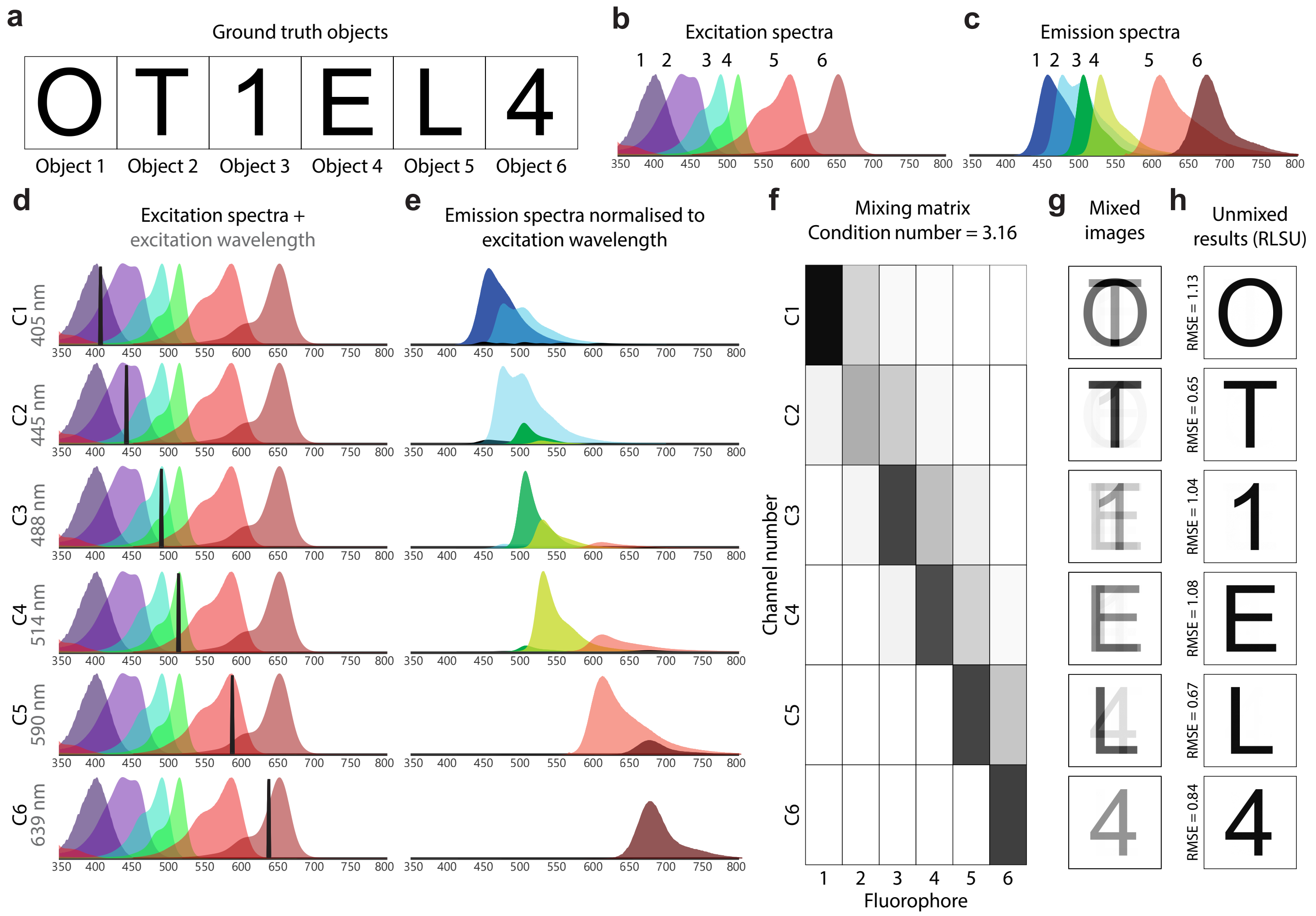


**Figure S1: a-h Simulations of excitation-based multispectral imaging approach.** **a** Simulated ground truth objects. **b** Simulated excitation spectra of six fluorophores used to label ground truth objects in (a). **c** Simulated emission spectra of six fluorophores used to label ground truth objects in (a). **d** Depiction of sequential laser excitation within each spectral channel (C) at different excitation wavelengths. **e** Emission spectra of each of the 6 fluorophores normalised to the excitation wavelength in each spectral channel. **f** Mixing matrix depicting the proportion of each fluorophore's fluorescence detected in each of the excitation channels. **g** Simulated spectrally-mixed, Poisson-noisy data. **h** Spectrally unmixed reconstructions of the ground truth objects using RLSU (1000 iterations).

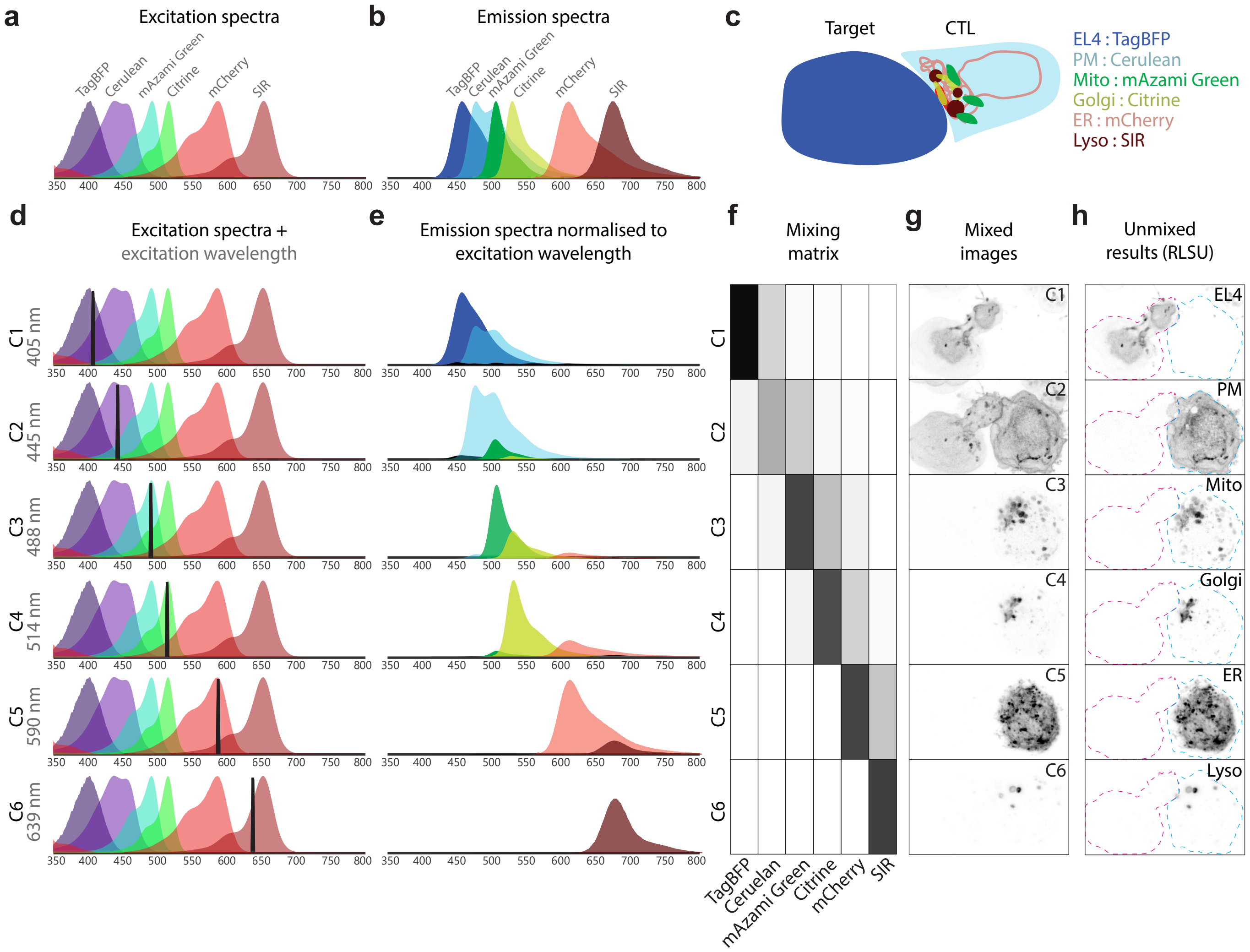


**Figure S2: a-h OT-1 CTLs were transfected with the ColorfulCell plasmid, stained with SiR Lysosome, and imaged during migration or 10 minutes after mixing with OVA-pulsed EL4 target cells. Individual Z-stacks were acquired using a sequential, excitation-based multispectral imaging pipeline followed by spectral unmixing using the RLSU algorithm (1000 iterations).** **a** Excitation spectra of six fluorophores (TagBFP, Cerulean, mAzami Green, Citrine, mCherry and SIR) used to label organelles in T cells and Target cells. **b** Corresponding emission spectra of six fluorophores (TagBFP, Cerulean, mAzami Green, Citrine, mCherry and SIR) used to label organelles in T cells and Target cells. **c** Graphical depiction of six fluorophores targeted to organelles in T cells and target cells **d** Depiction of sequential laser excitation within each spectral channel (C) at different excitation wavelengths **e** Emission spectra of each of the 6 fluorophores normalised to the excitation wavelength in each spectral channel. **f** Mixing matrix depicting the proportion of each fluorophore's fluorescence detected in each of the excitation channels. **g** Spectrally-mixed data acquired for each of the six excitation wavelengths using a LSM 980 Airyscan microscope (Zeiss) **h** Spectrally unmixed reconstructions of the underlying organelle structures using RLSU (1000 iterations). Dashed magenta line = target cell (OVA-pulsed EL4 cell) boundary; dashed cyan line = T cell boundary. EL4 = lymphoblastic target cell; PM = plasma membrane; Mito = mitochondria; Golgi = Golgi apparatus; ER = endoplasmic reticulum; lyso = lysosome.

| Figure | Modality | Cells | Pixel pitch (nm) | Z-spacing (nm) | $\lambda_{ex}$ (nm) | Labels | LUTs |
|---|---|---|---|---|---|---|---|
| Figure 1b,c<br>Figure 2a,c | Airyscan | OT-I | 30 | 130 | 405<br>445<br>488<br>514<br>590<br>639 | Lyn-Cerulean (PM)<br>MTS-mAzamiGreen (M)<br>GTS-Citrine (G)<br>mCherry-KDEL (ER)<br>SiR Lysosome (L) | Cyan<br>Red<br>Green<br>Orange<br>Magenta |
| Figure 1d-h<br>Figure 2b,c | Airyscan | OT-I<br>EL4 | 30 | 130 | 405<br>445<br>488<br>514<br>590<br>639 | Farnesyl-TagBFP (Target PM)<br>Lyn-Cerulean (PM)<br>MTS-mAzamiGreen (M)<br>GTS-Citrine (G)<br>mCherry-KDEL (ER)<br>SiR Lysosome (L) | Blue<br>Cyan<br>Red<br>Green<br>Orange<br>Magenta |
| Figure 3b | SDCM-M | OT-I<br>EL4 | 110 | 500 | 405<br>445<br>470<br>520<br>555<br>647 | Farnesyl-TagBFP (Target PM)<br>NLS-TagBFP (N)<br>Lyn-Cerulean (PM)<br>MTS-mAzamiGreen (M)<br>GTS-Citrine (G)<br>mCherry-KDEL (ER)<br>SiR Lysosome (L) | Duo-cyan<br>Blue<br>Grey<br>Red<br>Green<br>Circus-yellow<br>Magenta |
| Figure 3c | SDCM-M | OT-I<br>EL4 | 110 | 500 | 405<br>445<br>470<br>520<br>555<br>647 | Farnesyl-TagBFP (Target PM)<br>PACT-TagBFP (C)<br>NLS-TagBFP (N)<br>Lyn-Cerulean (PM)<br>MTS-mAzamiGreen (M)<br>GTS-Citrine (G)<br>mCherry-KDEL (ER)<br>SiR Lysosome (L) | Duo-cyan<br>Cyan<br>Blue<br>Grey<br>Red<br>Green<br>Circus-yellow<br>Magenta |
| Figure 3d | OPM | OT-I<br>EL4 | 111.6 | 500 | 405<br>488<br>561<br>638 | Farnesyl-TagBFP (Target PM)<br>NLS-TagBFP (N)<br>MTS-mAzamiGreen (M)<br>GTS-Citrine (G)<br>mCherry-KDEL (ER) | Duo-cyan<br>Blue<br>Red<br>Green<br>Circus-yellow |
| Figure 5a | SDCM | OT-I<br>EL4 | 103 | 500 | 405<br>488<br>561<br>640 | Farnesyl-TagBFP (Target PM)<br>CEPIA1er-eGFP (ER-Ca)<br>LifeAct-mApple (A)<br>SiR Lysosome (L) | Blue<br>Green<br>Magenta<br>Red |
| Figure 5c | SDCM | OT-I<br>EL4 | 103 | 500 | 405<br>488<br>561<br>640 | Farnesyl-TagBFP (Target PM)<br>MAPPER-eGFP (ER-PM)<br>LifeAct-mApple (A) | Blue<br>Green<br>Red |
| Figure 5e-f | SDCM-M | OT-I | 110 | 500 | 405<br>445<br>470<br>514<br>555<br>647 | PACT-TagBFP (C)<br>NLS-TagBFP (N)<br>Lyn-Cerulean (PM)<br>MTS-mAzamiGreen (M)<br>mCherry-KDEL (ER)<br>SiR Lysosome (L) | Grey (circle)<br>Blue<br>Cyan (dashed lin<br>Red<br>Circus-yellow<br>Magenta |

**Table 1: Experimental details and labelling information.** N, nucleus; PM, plasma membrane; M, mitochondria; G, Golgi; ER, endoplasmic reticulum; L, lysosomes; C, centrosome; A, actin; ER-PM, endoplasmic reticulum-plasma membrane contact sites; ER-Ca, endoplasmic reticulum calcium; SDCM-M, multispectral spinning disk confocal microscopy; SDCM, spinning disk confocal microscopy; OPM, multispectral oblique plane light sheet microscopy.

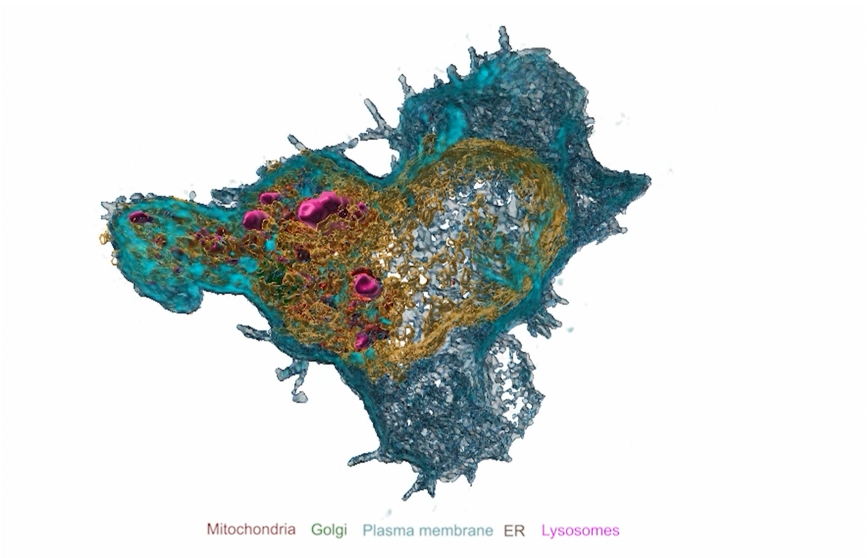


**Supplementary Video 1: Multispectral super-resolution imaging and 3D segmentation of organelles in a migrating CTL.**
Relates to Figure 1. Migrating CTL transiently expressing a polycistronic plasmid (Colorfulcell) co-stained with a live-cell dye (SiR lysosome) to simultaneously label lysosomes (magenta), mitochondria (red), plasma membrane (cyan), Golgi (green) and ER (orange) imaged using multispectral super-resolution followed by Airyscan processing and RLSU (1000 iterations). Processed, unmixed data presented before 3D segmentations of organelles to show organelle contacts and distributions in a representative migrating CTL ($n$ = 6 cells, $N$ = 3 independent experiments).

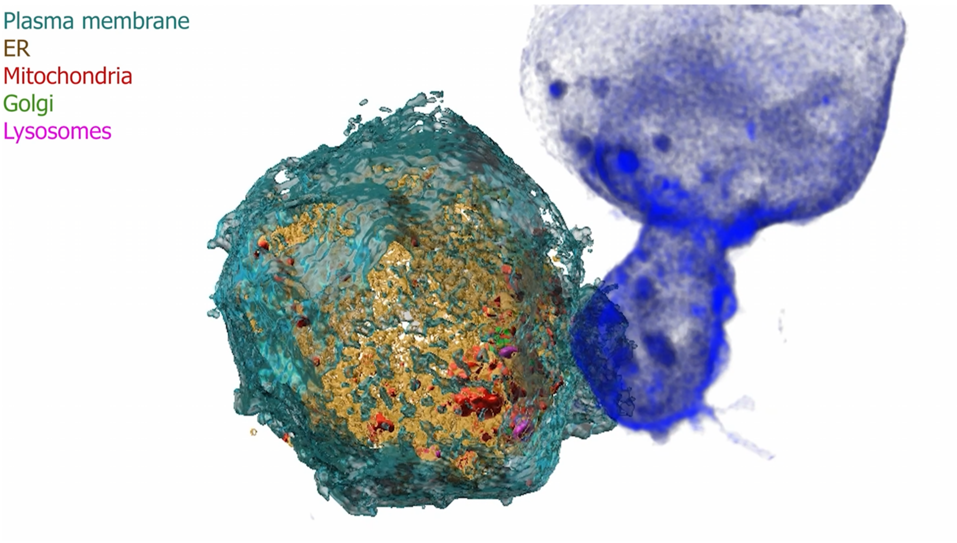


**Supplementary Video 2: Multispectral super-resolution imaging and 3D segmentation of organelles in a CTL during killing.**
Relates to Figure 1. A CTL transiently expressing a polycistronic plasmid (Colorfulcell) was co-stained with a live-cell dye (SiR lysosome) to simultaneously label lysosomes (magenta), mitochondria (red), plasma membrane (cyan), Golgi (green) and ER (orange) and imaged using multispectral super-resolution followed by Airyscan processing and RLSU (1000 iterations) after 10 minutes of target engagement. Processed, unmixed data presented before 3D segmentations of organelles to show organelle contacts and distributions in a representative CTL during killing ($n = 6$ cells, $N = 3$ independent experiments).

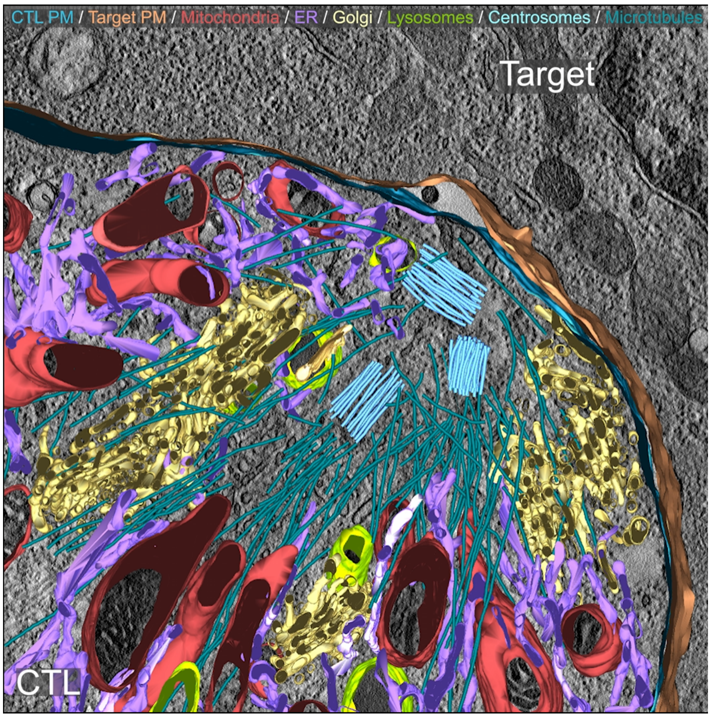


**Supplementary Video 3: 3D electron tomography of the immune synapse.**
Relates to Figure 2. Electron tomography (tilt series) followed by 3D segmentations shown of the CTL immune synapse during killing.

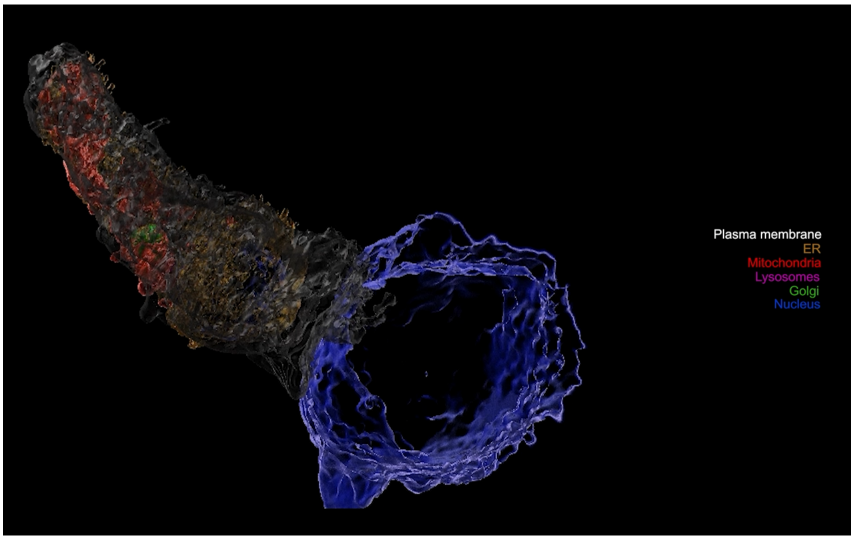


**Supplementary Video 4: Multispectral live-cell volumetric imaging of CTL killing using a multispectral spinning disk confocal microscope.**
Relates to Figure 3. A CTL transiently expressing a polycistronic plasmid (Colorfulcell) was co-stained with a live-cell dye (SiR lysosome) to simultaneously label nucleus (blue), lysosomes (magenta), mitochondria (red), plasma membrane (grey), Golgi (green) and ER (orange), with target-cell membrane in cyan and imaged using multispectral spinning disk confocal microscopy followed by deconvolution and RLSU (1000 iterations) during killing. Processed, unmixed data presented before 3D segmentations of organelles to show organelle contacts and distributions in a CTL during killing ($n$ = 6 cells, $N$ = 3 independent experiments).

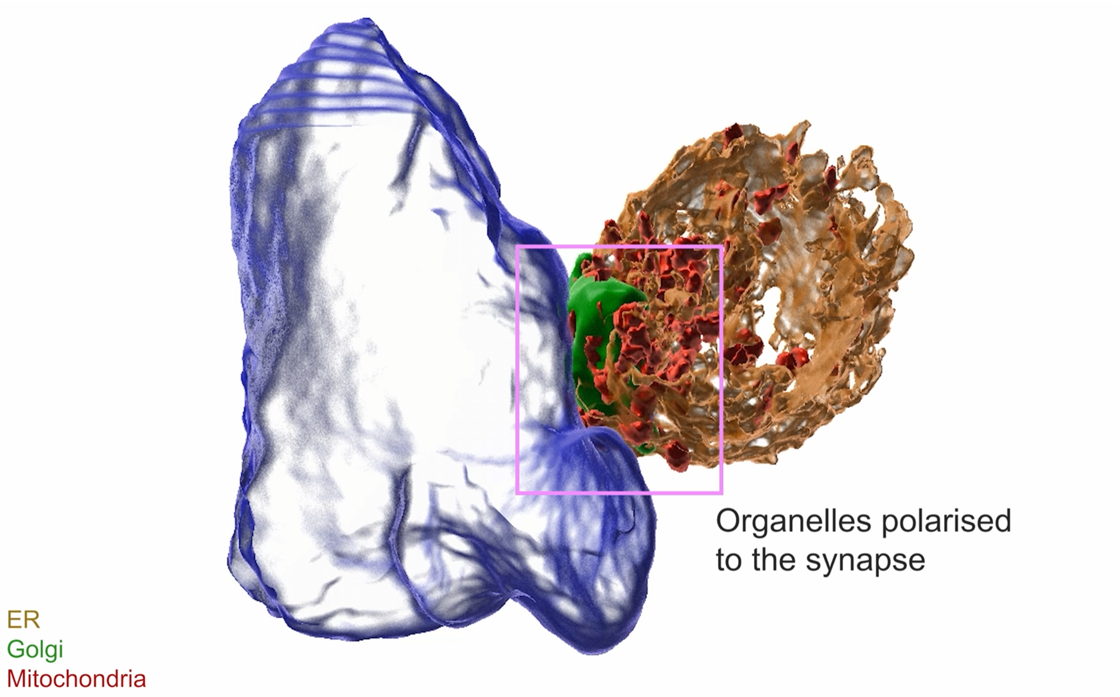


**Supplementary Video 5: Multispectral live-cell volumetric imaging of organelles polarised to the immune synapse using a multispectral oblique plane light sheet microscope.**
Relates to Figure 3. A CTL transiently expressing a polycistronic plasmid labelling mitochondria (red), Golgi (green) and ER (orange), imaged during organelle polarisation toward the target cell membrane (blue) acquired using multispectral oblique plane light sheet microscopy followed by RLSU (1000 iterations). Processed, unmixed data presented before 3D segmentations of organelles to show organelle contacts and distributions in a CTL during killing ($n = 6$ cells, $N = 3$ independent experiments).

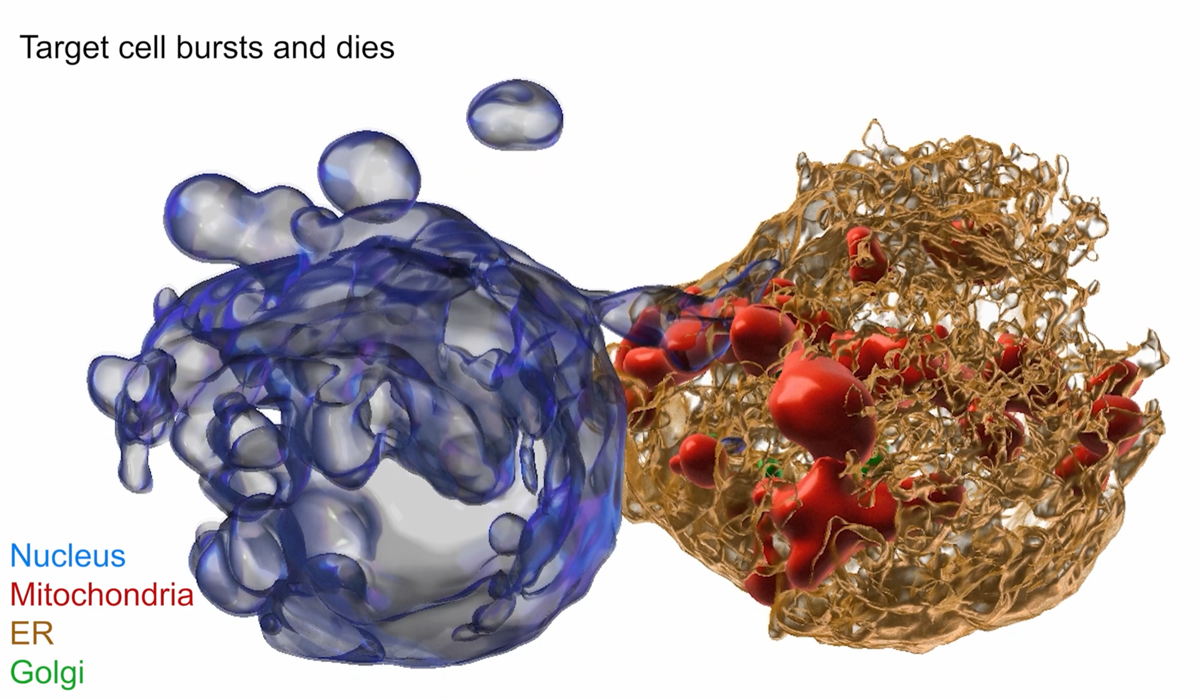


**Supplementary Video 6: Multispectral live-cell volumetric imaging of a CTL engaging and killing a target cell using a multispectral oblique plane light sheet microscope.**
Relates to Figure 3. A CTL transiently expressing a polycistronic plasmid labelling nucleus (blue), mitochondria (red), Golgi (green) and ER (orange), imaged during target killing (blue) acquired using multispectral oblique plane light sheet microscopy followed by RLSU (1000 iterations). Processed, unmixed data presented before 3D segmentations of organelles to show organelle contacts and distributions in a CTL during killing ($n = 6$ cells, $N = 3$ independent experiments). Note target death by the end of the movie.

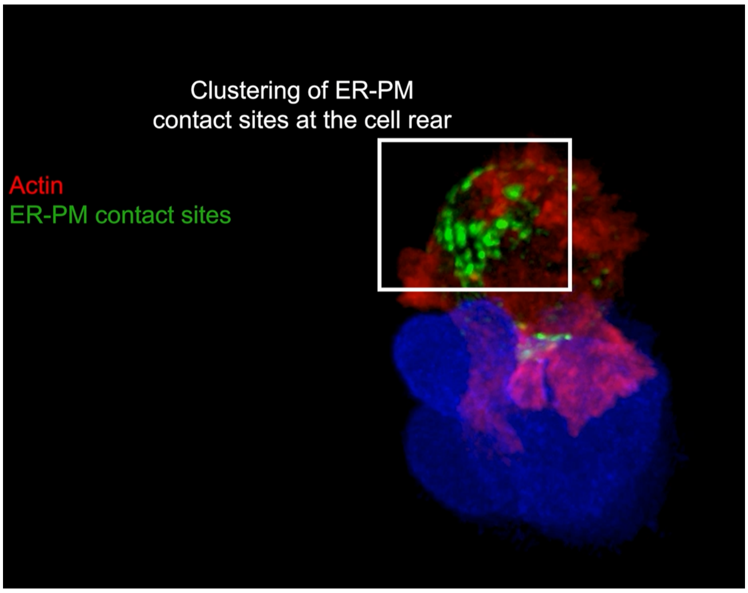


**Supplementary Video 7: Live-cell volumetric imaging of ER-PM contact sites and actin during CTL killing.**
Relates to Figure 5. A representative CTL ($n = 5$ cells, $N = 3$ independent experiments) transiently expressing MAPPER-GFP (ER-PM contact site reporter) and LifeAct-mApple was imaged live during killing.

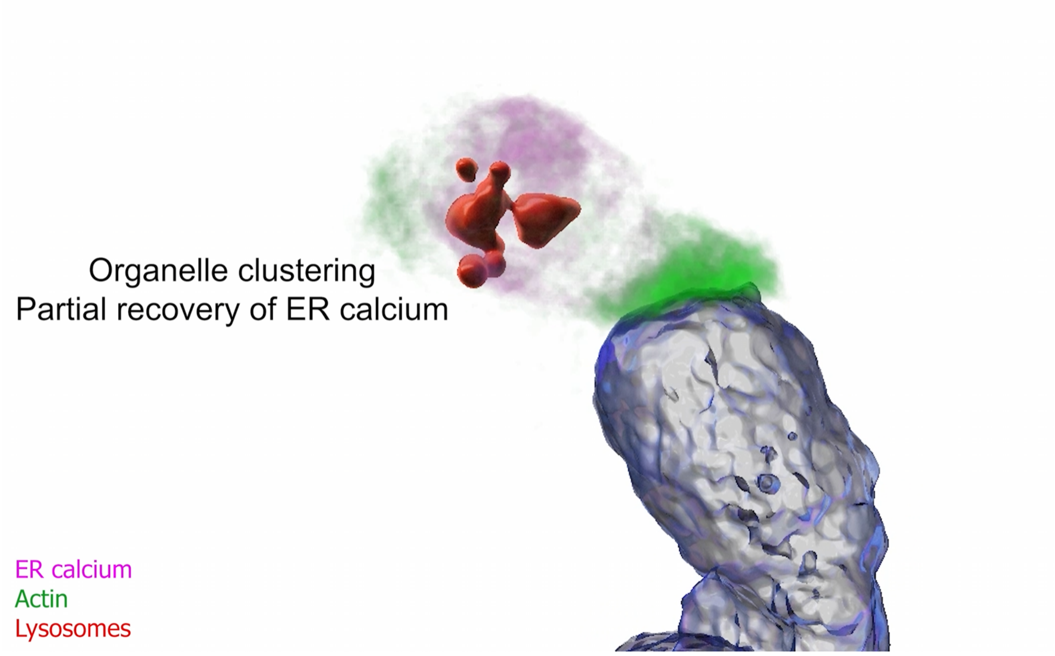


**Supplementary Video 8: Live-cell volumetric imaging of ER calcium, lysosomes and actin during CTL killing.**
Relates to Figure 5. A representative CTL ($n$ = 4 cells, $N$ = 3 independent experiments) transiently expressing CEPIA1er (ER lumenal calcium reporter) and LifeAct-mApple was co-stained with SiR lysosome and imaged live during killing.
